# Striking allelic diversity despite structural homogeneity of *ace-1* duplications in *Anopheles* mosquitoes

**DOI:** 10.1101/2023.05.10.539957

**Authors:** Jean-Loup Claret, Marion Di-Liegro, Alice Namias, Benoit Assogba, Patrick Makoundou, Alphonsine Koffi, Cédric Pennetier, Mylène Weill, Pascal Milesi, Pierrick Labbé

## Abstract

*A. gambiae s.l.* has been the target of intense insecticide treatment since the mid-XXth century to try and control malaria, and a substitution in the *ace-1* locus allowing resistance to organophosphate and carbamates insecticides has been rapidly selected for. Since then, several duplications of the ace-1 locus have been found in A. gambiae s.l. populations. They associate either several resistance copies (homogeneous duplications) or both resistance and susceptible copies (heterogeneous duplications). Heterogeneous duplications confer an intermediate trade-off between resistance in presence of insecticide and disadvantage in their absence. So far, and in striking contrast with *C. pipiens* mosquitoes, a single heterogeneous duplication had been describe in A. gambiae populations. We use an innovative approach, combining information from long and short read sequencing with Sanger sequencing to precisely identify and describe at least nine different heterogeneous duplications in A. gambiae. We further show that these alleles share the exact same structure than the previously identified heterogeneous and homogeneous duplications, namely 203-kb tandem amplifications with conserved breakpoints. Our study sheds a new light on the origin and maintenance of these alleles in A. gambiae populations, and pushes one step further the striking evolutionary convergence with *C. pipiens* mosquitoes.

## Introduction

Human activities have huge impacts on the environment, and these diverse anthropogenic modifications can lead to spectacular adaptations (see for example Otto 2018 or Hendry et al. 2017). Among those, resistance to biocides (e.g. antibiotic resistance in bacteria or resistance to pesticides in crop pests and disease vectors) are probably the most studied and the best understood, because of their crucial impacts on economy and public health. From an evolutionary biology point of view, these are also major models to understand the dynamics of adaptations from their genesis, *i.e.* deciphering the genetics of the adaptation (e.g. polygenic *vs* mono-or oligo-genic) or to understand how the evolutionary processes shape these dynamics (e.g. spatial variation in selective pressure intensity, fluctuating selection overtime; Guillemaud *et al*. 1998, David *et al*. 2010, Milesi *et al*. 2016). Studying these resistances notably revealed the complexity and the diversity of the genomic structural rearrangements underlying adaptations, well beyond the role of single nucleotide polymorphisms (SNPs). For example, cases of xenobiotic resistances linked to gene duplications are plenty (Devonshire & Sawicki, 1979; Leister *et al*. 2004; Labbé *et al*. 2007; Kwon *et al*. 2010; Patterson *et al*. 2014). In the present study, we focused on one of those well-known models, the case of insecticide resistance in the malaria-vector mosquito *Anopheles gambiae s.l.*, and in particular on the genomic nature and diversity of the resistance alleles of the *ace-1* locus.

*A. gambiae s.l.* has indeed been the target of intense insecticide treatment since the mid-XX^th^ century, particularly on the African continent, to try and control malaria (619,000 deaths in 2021, see https://www.who.int/news-room/fact-sheets/detail/malaria). While pyrethroids (PYR) are the most commonly-used insecticides, organophosphate (OP) and carbamates (CX) have also been used since the 50’s. They target the acetylcholinesterase (AChE), an enzyme that regulates the activity of the synaptic neurotransmitter acetylcholine (Weill *et al*. 2003). A substitution in the AChE-encoding gene *ace-1*, resulting in a glycine to serine substitution at the 280 codon of the protein (G280S), allows resistance to OP and CX by hindering their binding to AChE (R allele, Weill *et al*. 2004, also referred to as G119S mutation according to its position in the homolog gene of *Torpedo californica* where AChE structure was first studied). This mutation has been selected for, independently, in multiple mosquitoes species (Huchard *et al*. 2006; Weill *et al*. 2003). However, the enzymatic activity of the protein encoded by the R allele is 60% lower than that of its wild-type susceptible counterpart (S allele; Bourguet *et al*. 1997, Alout *et al*. 2008, Labbé *et al*. 2014); as a result, the R alleles are strongly selected against in the absence of insecticide.

Other types of mutations have also been selected in response to the use of these insecticides, including structural variants (SVs). In particular, several duplications of the *ace-1* locus have been found in *A. gambiae s.l.* (Fig. 1A), associating either several R copies, *i.e.* homogeneous duplications (R^x^ alleles; Assogba et al 2016; Grau-Bové 2021), or both R and S copies, *i.e.* heterogeneous duplications (D alleles; Assogba 2015, 2016; Grau-Bové 2021). R^x^ alleles confer higher resistance levels and are therefore favored in highly-treated areas. However, the heterogeneous duplications (D alleles) allow the fixation of the heterozygous phenotype (Labbé et al. 2014, Assogba et al. 2015), *i.e.* an intermediate trade-off between resistance in presence of insecticide and disadvantage in their absence (aka “selective cost”, but see Lenormand et al. 2018). Hence, D allele are more advantageous in populations exposed to intermediate selective pressures, either because of reduced concentration of treatments *per se*, or because of temporal or geographical variations in treatment intensity (Labbé et al. 2014, Milesi et al. 2017).

**Figure 1.**
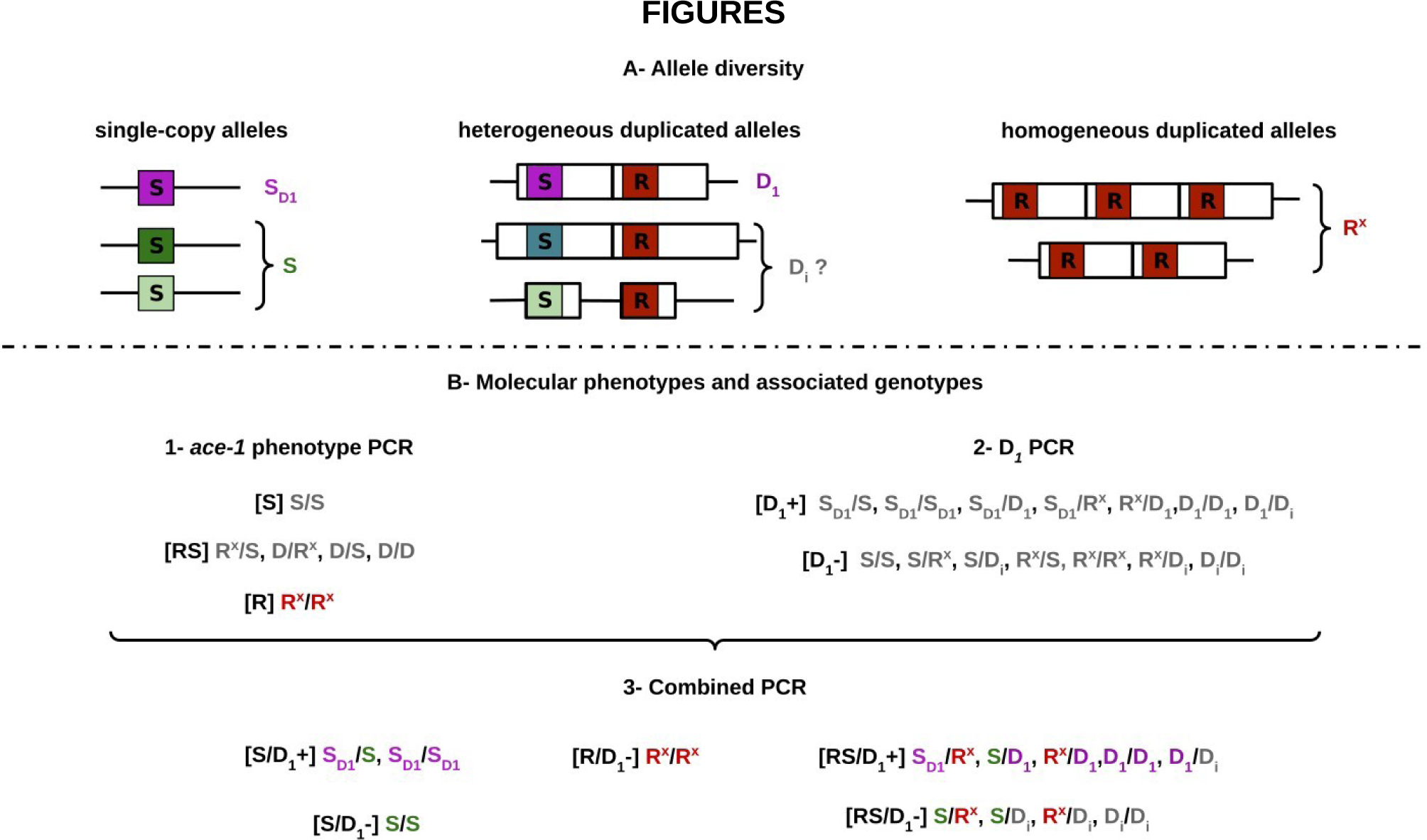
Diversity of *ace-1* alleles, and molecular phenotyping. (Modified from Assogba *et al*. 2018). The top part of the figure (A) illustrates the various alleles found at the *ace-1* locus: different single-copy S alleles on the left (green) and homogeneous duplicated alleles R^x^ on the right (here with 2 or 3 R copies, R^3^ and R^2^ resp., in red). The central part illustrates the known D_1_ heterogenous allele, with its D(S) copy (in green) and its D(R) copy (in red), as well as putative architectures for other heterogeneous D_i_ alleles (depending on the size of the amplified region). NB: the single-copy S_D1_ allele has the same sequence than the D(S) copy of *ace-1* (hence the same color). The bottom part (B) illustrates the two PCR used to identify the genotypes of triple peaks samples. The combined information of the « *ace-1* phenotype » PCR (1) and the « D_1_ PCR » (2) allow the partial discrimination of 5 phenotypes with 13 possible genotypes (3).

Diversity in duplicated alleles can result from two types of variations, i) variation in the DNA sequence of the amplicons, in particular the *ace-1* sequence, and/or ii) variation in the number of copies. In the present study, we will mostly focus on the first type of variants, but we will consider both kinds to be different *ace-1* alleles. We will thus refer to alleles carrying different *ace-1* sequences as “sequence-alleles”, and those differing in numbers of copies as “copy-number-alleles”. In *A. gambiae s.l.*, the same R sequence-allele has been found in several African countries, in R^x^ alleles, *i.e.* with several copy-number-alleles (Assogba 2016). Only one D allele, named Ag-D_1_ (thereafter D_1_ for simplicity), has been formally described, using direct sequencing of cloned fragments of the *ace-1* gene (*i.e.* real haplotypes, Djogbénou *et al*. 2008). D_1_ carries two *ace-1* copies, one R copy (identical to that found in R^x^ alleles) and one S copy, in 203 Kb tandem amplicons (Assogba *et al*. 2016). D_1_ is found all over West Africa (Assogba *et al*. 2018), which is in sharp contrast with the 27 D sequence-alleles found in *C. pipiens*, with several different D sequence-alleles segregating in a same population (Milesi *et al*. 2018). However, Grau-Bové et al. (2021) recently analysed a large dataset of Illumina paired-end genomes of *A. gambiae s.l.* from all over Africa (The *Anopheles gambiae* 1000 Genomes Consortium (2021): Ag1000G phase 3).

Based on the variations in depth of coverage of the alternative bases at position 119 (*i.e.* of R or S sequences), they suggested that several copy-number-alleles, differing in their numbers of R and S copies, might actually be segregating in Africa. The nature of their data (*i.e.* short read sequencing) made it hazardous to assess whether several D sequence-alleles were segregating in these populations and a fortiori to identify potential new sequences-alleles.

Evidencing the existence of a diversity of D alleles, whether copy-number-alleles or sequence-alleles, could help us understand the origin of these mutations, and more generally how SVs are selected for short-term adaptation. We thus tried a new approach to assess the sequence diversity of *ace-1* alleles, and the number of R and S copies in heterogeneous duplications, and used information from different inputs to understand the origin of this diversity. As previous studies have shown that the frequency of D alleles in *A. gambiae s.l.* was particularly high in Ivory Coast (Assogba et al. 2018), and as Grau-Bové et al. (2021) suggested that several D copy-number-alleles could segregate there, we analysed the structures and diversity of the heterogeneous duplications present in two natural populations of Ivory Coast, Yamoussoukro and Yopougon. By screening samples from Assogba et al. (2018) for the presence of D_1_ allele, we found that it could not explain alone the observed frequencies of D alleles. By sequencing a large part of the *ace-1* locus for several individuals, we obtained the various haplotypes of each of their copies through both traditional cloning and a more recently-developed approach based on long-read sequences of PCR products (Namias *et al*. 2022). We thus revealed that at least nine different *ace-1* D alleles segregate in these two populations. Using whole genome sequencing, we then showed that at least 5 of these alleles share the exact same structure, *i.e.* two 203kb tandem amplicons with the exact same breakpoints. Finally, we discuss what these findings suggest in terms of duplication genesis, but also on the role of SVs in the adaptation process.

## Material & Methods

### Sampling and identication

We focused on two localities from Ivory Coast where the presence of heterogeneous duplications have already been documented (Assogba *et al*. 2018, Grau-Bové *et al*. 2021), and for which resistance has been monitored since 2012. We used preserved DNA extracted from samples collected in 2012, 2015, 2016 (Assogba et al. 2018) and we collected new samples in 2019. They were identified as *A. coluzzii* through a multiple PCR protocol: a first PCR allows the discrimination of *A. arabiensis* from *A. gambiae s.s.* and *A. coluzzii* (Supporting information Tab. 1, “Species”; Scott *et al*. 1993), and a second one distinguishes *A. gambiae s.s.* from *A. coluzzii* (Supp. info. Tab. 1, “Form”; Favia *et al*. 1997). We completed the dataset with publicly available sequence data of the *ace-1* gene from samples collected at the same period in the same or nearby countries (Ghana, sequences from Weetman *et al*. 2015; and Benin, Burkina Faso and Ivory Coast from Djogbenou *et al*. 2008; see Supp. info. Tab. 2).

### DNA extraction and PCR conditions

We extracted DNA from individual L4 following a protocol modified from Collins *et al*. (1987). Briefly, each larva was ground in 200µl CTAB buffer (100mM Tris HCL, pH8.0, 10mM EDTA,1.4M NaCl, 2% CTAB), then incubated for 15 minutes at 60°C. 200µl of chloroform with 4% of isoanyl alcohol were added, and the solution was centrifuged for 10 minutes at 8000 rotations/min. The supernatant was transferred to a new tube with 200µl of isopropanol to precipitate DNA at room temperature. DNA was washed with 400 µl of 70% ethanol after 10 minutes of centrifugation (10000 rotations/min), dried, and then rehydrated in 50µl H_2_O.

All the PCR described below were realized using the Promega PCR kit (Madison, Winsconsin, USA) with *ca*. 50 ng of genomic DNA into 40 μL of PCR-mix and using the following: 94°C for 30s, annealing temperature for 30s, and 72°C for 1 to 2 min for a total of 30 cycles (primers and annealing temperatures are listed in Supp. info. Tab. 1).

### Heterogeneous duplication detection and frequency estimation

#### ace-1 phenotyping

We performed the “*ace-1* phenotype” PCR-RLP test described in Djogbénou et al. (2008): it amplifies a 817 bp sequence of the *ace-1* locus encompassing the resistance-diagnostic G119S mutation. This mutation generates an *AluI* restriction site and allows the distinction between three phenotypes: resistant homozygous [RR], susceptible homozygous [SS], and heterozygous [RS] (Fig. 1). However, it does not allow the differentiation between standard heterozygous individuals for single-copy alleles (RS) and individuals carrying a heterogeneous duplicated allele (D) (Assogba et al. 2015), as D alleles associate both susceptible D(S) and resistant D(R) copies of the *ace-1* locus.

#### Estimation of D allele frequencies

D allele frequencies can nevertheless be inferred from the phenotypic frequencies (Tab. 1), as their presence in a population causes an excess of [RS] relatively to what is expected under Hardy-Weinberg equilibrium (Lenormand *et al*. 1998). We thus took advantage of this observation and used the same approach as in Assogba *et al*. (2018) to compute the D allele frequencies, independently for each year and location, implementing the maximum-likelihood approach developed by Lenormand et al. (1998). We calculated the log-likelihood, *L*, of observing all the data as follow:

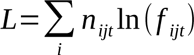

**Table 1:** *ace-1* phenotype diversity in Yamoussoukro and Yopougon. Samples (identified by year and locality) originate from Assogba et al.’s (2018) study. [RR], [RS] and [SS] phenotypes were obtained through the “*ace-1* phenotype” PCR (see Material and Methods, Fig. 1B). The number of [D_1_+] individuals are indicated in brackets (“D_1_” PCR, Fig. 1B).

| Locality | Year | Resistance phenotype |  |  |  |
| --- | --- | --- | --- | --- | --- |
|  |  | [RR] | [SS] ([D<sub>1</sub>⁺]) | [RS] ([D<sub>1</sub>⁺]) | N |
| Yopougon | 2012 | 0 | 40 (1) | 15 (3) | 55 |
|  | 2015 | 1 | 13 (0) | 45 (25) | 59 |
|  | 2016 | 1 | 20 (2) | 37 (18) | 58 |
| Yamoussoukro | 2012 | 0 | 21 (1) | 25 (14) | 46 |
|  | 2015 | 1 | 12 (2) | 35 (15) | 48 |
|  | 2016 | 0 | 9 (1) | 30 (15) | 39 |

with n_ijt_ and f_ijt_, the observed number and the predicted frequency of individuals with phenotype *i* in population *j* at time *t*, respectively. It was maximized (L_max_) for each sample using a simulated annealing algorithm (Labbé et al., 2009; Lenormand, et al., 1998). The support limits (SL, equivalent to 95% confidence intervals) were defined as the minimum and maximum allele frequencies that did not significantly decrease the likelihood. Recursions and likelihood maximization algorithms were written and compiled with Lazarus v1.0.10 (http://www.lazarus.freepascal.org/).

#### Discriminating new D alleles from D_1_ allele and standard heterozygotes

Before this study a single D allele had been characterized in *A. gambiae*, referred to as D_1_ (Assogba *et al*. 2018, Grau Bové *et al*. 2021). For each population and year, we thus tested whether this allele alone could explain the estimated frequency of D alleles, or if more alleles (hereafter D_i_) could segregate in the populations.

To do so, we used a PCR-RFLP test specific to the D_1_ susceptible *ace-1* copy (“D_1_” PCR-RFLP-test, Supp. mat. 1; Assogba et al. 2015) on all individuals with a [RS] phenotype (*i.e.* those that could harbor a D allele): a 817 bp fragment of the *ace-1* gene is PCR-amplified using specific primers and an *AvaI* restriction site specific to the D_1_(S) copy allows distinguishing between D_1_ carriers ([D_1_+] phenotype) and individuals that do not carry D_1_ ([D_1_-] phenotype; Fig. 1). We then compared a model considering only three alleles (R, S, D_1_) with models considering either four (R, S, S_D1_, D_1_) or five alleles (R, S, S_D1_, D_1_, D_i_), using likelihood ratio tests (see Labbé et al., 2009; Milesi et al., 2016); we thereby took into consideration the possibility of occurrence of single-copy susceptible alleles S_D1_, carrying the same *AvaI*-diagnostic mutation as the D_1_(S) copy.

As our goal was to characterize the sequences of new D_i_ alleles, we first sequenced (Sanger sequencing ABI 3500 xL, Applied Biosystems by Thermo Fisher Scientific) the *ace-1* PCR product for all the [RS] individuals that were [D1-] (*i.e.* not amplified by the PCR-RFLP test specific to the D_1_ allele), to prevent the presence of D_1_ (except for a few controls). If these individuals harbor at least one D_i_ allele (genotypes D_i_R, D_i_S or D_i_D_j_), then at least three copies of the *ace-1* locus should be present in the PCR product. Providing that the different copies carry different single nucleotide polymorphisms (SNPs), one could expect diagnostic “triple peaks” (*i.e.* positions at which three different SNPs can be found) in this mix sequence. This allows discriminating D_i_ carriers from standard RS heterozygotes. Although powerful to detect new D alleles, this approach can lead to an underestimation of the new D_i_ allele frequency, as *ace-1* copies can be similar between D alleles and single-copy resistance or susceptible alleles, so that “triple peaks” are not detected.

### Heterogeneous duplication diversity

#### TA cloning

To identify the different *ace-1* haplotypes found in individuals identified as carrying a new D_i_ allele, we purified the products of the *ace-1* PCR (see above) using the BS664-250 Preps EZ-10 Spin Column PCR Purification kit (New England BioLabs, Evry France). The purified PCR products were then cloned (TOPO TA Cloning Kit pCR 2.1-TOPO Vector and TOP10F’ invitrogen bacteria). For each individual, we genotyped 24 clones (Supp. Info. Tab. 3) using the *AluI* RFLP test (to discriminate R and S copies). All R sequences currently known in *A. coluzzii* and *A. gambiae* are identical on this 817pb fragment, so most of the diversity is expected to result from variation between D(S) copies (in *C. pipiens*, the 27 different D alleles described so far mostly differ by their D(S) copies, Milesi et al. 2018). We thus sequenced one resistance copy (*i.e.* D(R) or R), and at least 11 susceptible copies (*i.e.* D(S) or S; ABI 3500 xL, Applied Biosystems by Thermo Fisher Scientific).

#### Nanopore sequencing of ace-1 PCR products

It has been shown that Nanopore sequencing of PCR products can be used to reconstruct the haplotypes of multi-copy gene families, in spite of Nanopore sequencing errors, a method less tedious and more sensitive than TA cloning (Namias et al, 2022). We thus directly sequenced the purified *ace-1* PCR product of 12 [RS, D_1_-] individuals displaying triple peaks (Supp. Info. Tab. 3) using Nanopore long-reads technology to capture, in a single read, each full 817 bp amplicon (5 individuals had been previously analysed with TA cloning to assess the reliability of the Nanopore sequencing-based approach). We then adapted the method developed by Namias *et al*. (2022): reads were mapped on a reference file containing two reference sequences, one R and one S, using *minimap2 v.2.24* (Li, 2018) with the options *map-ont* and without secondary alignments. SNPs were then called using *bcftools 1.15*, with the *config*-*ont* option, and a minimum mapping quality of 10. Finally, haplotype phasing was performed using *WhatsHap 1.4* (Martin et al, BiorXiv 2016) with the default options. As mentioned in Namias *et al*. (2022), some heterozygous SNPs on the S sequences were not called, although they were supported by a high number of reads, because the read distribution did not fit with a diploid framework (with only two S copies). We used the script provided by Namias *et al*. (2022) to recover those SNPs.

#### ace-1 sequence trees

To assess the diversity of *ace-1* alleles obtained either through TA cloning or Nanopore sequencing (one R and different S), we aligned all these sequences and compared them using a Neighbour-Joining phylogram compiled by *MEGA* software (MEGA11: Molecular Evolutionary Genetics Analysis version 11; Tamura, Stecher, and Kumar 2021). We added to this phylogram several known *ace-1* sequences: a R reference sequence and the reference D_1_(S) copy. We used the information from molecular tests, whole genome sequencing, *ace-1* PCR product long read sequencing and sequence frequency to discriminate D(S) copy from S allele for each individual.

To infer the geographical origin of the newly characterized D alleles, we then added S alleles described in previous studies and sampled from 2012 to 2019 in Ivory Coast, Benin, Burkina Faso and Ghana natural populations (Weetman *et al*. 2015; Assogba *et al*. 2018, see Supp. Info. Tab. 2) to our alignment. We then computed a maximum likelihood phylogram using a Tamura 3 parameters model with gamma distributed with invariant sites (G+I) mutation rates (best model fit determined with MEGA11). The phylogram was then plotted using the *ggplot2* (Wickham H. 2016) and *ggtree* package (Yu G. 2022) in *R*.

### Duplication architecture

#### Genomic characterisation

To determine the structural architecture of the newly characterized D_i_ alleles, we followed the protocol developed by Assogba *et al*. (2016) for D_1_. Twelve individuals sampled in Yopougon and Yamoussoukro (Supp. Info. Tab. 3), carrying putative new duplications (identified using the protocols described above) were sequenced (Illumina paired-end sequencing, 150pb reads, 350 pb insert size). Reads from each individual were mapped to the *A*. *gambiae* PEST reference genome assembly (AgamP4.13; https://www.vectorbase.org) using the *bwa* (*-mem*) algorithm (Li & Durbin, 2009). The per-base depth of coverage (*pb*DoC) of the region surrounding *ace-1* (from 2Mb to 5Mb) on the 2R chromosomal arm) was obtained using the *samtools* suite (Danecek *et al*. 2021). To plot the *pb*DoC along this chromosome, we standardized them (*pb*DoC*_std_*) by dividing them by the average *µ*DoC calculated over the whole 2R chromosome (*pb*DoC*_std_* = *pb*DoC/*µ*DoC), using the R software (v.4.2.2, R Core Team, 2022; https://www.R-project.org/). It allowed a fine scale observation of the structure of the duplications (location, size, gene copy number, etc.). We further combined information from insert-size and local enrichment in soft-clipped reads to precisely map the breakpoints of the duplication.

#### Molecular validation of structural homologies

Assogba et al. (2016) developed a diagnostic PCR test for R^x^ and D_1_ duplications, which amplifies a 460bp sequence overlapping the junction between the amplicons (Supp. info. Tab. 1, “Junction”). We used this PCR to further assess whether the newly identified D_i_ alleles also shared this junction, and thus the 3’ breakpoint of the D_1_ and R^x^ alleles, using some susceptible (SS) individuals as controls.

## Results

### Ivory Coast populations are highly polymorphic for D alleles

We first analysed samples collected in two populations of Ivory Coast (Yamoussoukro and Yopougon) for which resistance has been monitored from 2012 to 2016 (Assogba *et al*. 2018) (Tab. 1). We built a first model (Model A) to estimate the frequencies of three alleles (R, S and D) from the [SS], [RS] and [RR] phenotypes (“*ace-1* phenotype” test): the two populations showed a significant excess of heterozygotes in comparison with expectations under Hardy-Weinberg equilibrium when considering only R and S alleles (LRT, 9.27 > X^2^ > 24.42, *p-value* > 0.01**, except for 2012 in Yopougon), thus suggesting the presence of D alleles at relatively high frequencies (>0.15; Tab. 2A).

**Table 2:** Estimated allele frequencies under different models. For each locality and sampling year, the frequencies of the different alleles (along with their support limits, *i.e.* roughly equivalent to 95% confidence intervals, brackets) have been estimated using a maximum likelihood approach under different assumptions (see text). Model A - The first model considers only the phenotypes resulting from the “*ace-1* phenotype” test, thus 3 alleles, R, S and D, for 3 phenotypes and 6 genotypes (Fig. 1B). LRT corresponds to the likelihood ratio test between this model and another one without any D allele (chi-square distribution with 1 DDL). Model B **-** The second model considers the phenotypes resulting from the combination of the “D_1_” and “resistance phenotype” tests, *i.e.* adding the specific information on D_1_ frequency. Four alleles are thus considered, R, S, S_D1_ (as some [SS] are also [D_1_+]) and D_1_, for 5 phenotypes and 10 genotypes (Fig. 1B). Model C-The third model analyzes the same data, but considers 5 alleles, R, S, S_D1_, D_1_ and D_i_ (*i.e.* at least another D allele), for 5 phenotypes and 15 genotypes (Fig. 1B). The *p*-value of the LRT comparing models B and C (chi-square distribution with 1 DDL) is indicated for each population.

| Model A- 3 alleles model |  |  |  |  |  |  |  |  |
| --- | --- | --- | --- | --- | --- | --- | --- | --- |
| Locality | Year | *R* | *S* | *D* | LRT ($\chi^2$) | *p*-value | | |
| Yopougon | 2012 | 0.00 [0.00:0.18] | 0.85 | 0.15 [0.00:0.22] | 2.38 | 0.12<sup>NS</sup> |  |  |
|  | 2015 | 0.13 [0.03:0.27] | 0.47 | 0.40 [0.23:0.55] | 24.42 | 7.73x10<sup>-7***</sup> |  |  |
|  | 2016 | 0.13 [0.03:0.27] | 0.61 | 0.27 [0.11:0.41] | 11.94 | 5.50x10<sup>-4***</sup> |  |  |
| Yamoussoukro | 2012 | 0.00 [0.00:0.20] | 0.68 | 0.32 [0.10:0.43] | 9.27 | 2.33x10<sup>-3**</sup> |  |  |
|  | 2015 | 0.14 [0.03:0.29] | 0.53 | 0.33 [0.16:0.49] | 14.84 | 1.17x10<sup>-4***</sup> |  |  |
|  | 2016 | 0.00 [0.00:0.22] | 0.50 | 0.50 [0.26:0.63] | 19.27 | 1.13x10<sup>-4***</sup> |  |  |
| Model B- 4 alleles model |  |  |  |  |  |  |  |  |
| Locality | Year | *R* | *S* | *S*<sub>D1</sub> | *D*<sub>1</sub> |  |  |  |
| Yopougon | 2012 | 0.11 [0.06:0.18] | 0.85 | 0.01 [0.00:0.05] | 0.03 [0.01:0.07] |  |  |  |
|  | 2015 | 0.25 [0.16:0.34] | 0.51 | 0.00 [0.00:0.03] | 0.24 [0.16:0.33] |  |  |  |
|  | 2016 | 0.21 [0.14:0.30] | 0.60 | 0.03 [0.00:0.08] | 0.16 [0.09:0.24] |  |  |  |
| Yamoussoukro | 2012 | 0.14 [0.08:0.23] | 0.68 | 0.02 [0.00:0.07] | 0.16 [0.09:0.25] |  |  |  |
|  | 2015 | 0.27 [0.18:0.38] | 0.54 | 0.04 [0.01:0.11] | 0.15 [0.08:0.24] |  |  |  |
|  | 2016 | 0.24 [0.15:0.36] | 0.53 | 0.02 [0.00:0.10] | 0.20 [0.11:0.31] |  |  |  |
| Model C- 5 alleles model |  |  |  |  |  |  |  |  |
| Locality | Year | *R* | *S* | *S*<sub>D1</sub> | *D*<sub>1</sub> | *D*<sub>i</sub> | LRT ($\chi^2$) | *p*-value |
| Yopougon | 2012 | 0.00 [0.00:0.17] | 0.84 | 0.01 [0.00:0.05] | 0.03 [0.00:0.07] | 0.12 [0.00:0.20] | 1.60 | 0.21<sup>NS</sup> |
|  | 2015 | 0.13 [0.03:0.27] | 0.47 | 0.00 [0.00:0.04] | 0.24 [0.16:0.33] | 0.16 [0.01:0.31] | 4.61 | 0.03* |
|  | 2016 | 0.13 [0.03:0.27] | 0.58 | 0.03 [0.00:0.09] | 0.16 [0.09:0.24] | 0.11 [0.00:0.25] | 2.37 | 0.12<sup>NS</sup> |
| Yamoussoukro | 2012 | 0.00 [0.00:0.20] | 0.67 | 0.02 [0.00:0.07] | 0.16 [0.09:0.25] | 0.16 [0.00:0.26] | 2.30 | 0.13<sup>NS</sup> |
|  | 2015 | 0.14 [0.03:0.29] | 0.49 | 0.04 [0.01:0.12] | 0.15 [0.08:0.24] | 0.18 [0.02:0.35] | 4.99 | 0.03* |
|  | 2016 | 0.00 [0.00:0.22] | 0.47 | 0.03 [0.00:0.11] | 0.20 [0.11:0.31] | 0.30 [0.07:0.44] | 7.07 | 0.01** |

To test whether D_1_ alone could explain the D allele frequencies estimated above, we screened all [RS] individuals (and a few [SS] as controls), using a molecular test specific to D_1_(S), the S copy of the D_1_ allele (“D_1_” test; Assogba et al. 2015). It turned out that some rare [SS] individuals were also positive for the test (Tab. 1), suggesting that at least one S allele (thereafter named S_D1_) shares the same diagnostic mutation as D_1_(S) and segregates in these populations. We thus built a second model (Model B) considering four alleles (R, S, S_D1_ and D_1_ only), to analyze the phenotypes resulting from the combination of the “*ace-1* phenotype” and “D_1_” tests (Fig. 1B). However, the frequencies for the R allele appeared much higher than expected, while the frequency of D_1_ were much lower than those found in Model A (Tab. 2B, Tab. 1). Suspecting that this could result from the presence of other D alleles (D_i_), we thus compared model B with a third model considering five alleles (R, S, S_D1_, D_1_ and D_i_). Model C fitted significantly better than model B for three out of the six populations (Yopougon 2015, and Yamoussoukro 2015 and 2016, *p* <0.05, Tab. 2C), with more coherent R and overall D frequencies (Tab. 2A and C *vs* B), strongly supporting the presence of at least one other D allele segregating in these populations. Interestingly, while the yearly sums of the frequencies of D_1_ and D_i_ in model C are similar to that of the overall D allele frequencies estimated in model A (Tab. 2), D_1_ was not always the most frequent D allele (e.g. Yamoussoukro 2015 and 2016).

We first wanted to sequence only D alleles that were different from D_1_: we thus sequenced a 817 bp PCR fragment of the *ace-1* locus for all the [RS, D_1_-] individuals, *i.e.* individuals that could carry a D allele but not D_1_ (N = 97, Tab. 1), as well as three individuals with D_1_ as controls. The Sanger sequences of the *ace-1* PCR products were a mix of R and S copies. Some individuals (N = 22) displayed several positions with triple peaks (thereby confirming the existence of at least three sequences in the mix), and, as they are negative for D_1_ specific features, it confirms the presence of other D alleles (D_i_) segregating in *A. gambiae s.l.* populations.

In order to assess whether the new D alleles could still be present at the time of the study, and to get a hint of their current frequencies, we also sequenced all [RS] individuals found in samples collected in 2019 in Yopougon and Yamoussoukro (N= 27 for each). We found 6 more individuals whose electrophoregram displayed triple peaks, 5 of which, were compatible with D_1_. From 2012 to 2019, we thus recovered a total of 28 “triple-peak” individuals.

To describe the diversity of the D_i_ alleles, the PCR product of 22 of these 28 individuals were cloned (Supp. Info. Tab. 3), and at least 10 clones were sequenced per individual. As expected, for each individual, three different *ace-1* sequences were identified, one with the G119S mutation providing the resistance (R), and two susceptible copies but with discriminating mutations.

This cloning and sequencing protocol is however tedious, and thus difficult to apply to large numbers of individuals. We therefore tested whether another approach that had been developed for *Wolbachia Cid* genes multigenic family (Namias *et al*. 2022) could be used here: the PCR product (*ace-1*, 817 bp fragments) of twelve individuals was thus directly sequenced using Nanopore long-reads. Six individuals among the 22 cloned, and for which the sequences of the *ace-1* copies were thus known (among which some carried D_1_), served as positive controls; we also analysed the last six of the 28 “triple-peak” individuals using this long-read-based approach (Supp. Info. Tab. 3). Using a specifically-designed bioinformatics pipeline, we were able to clearly identify three sequences of the *ace-1* locus for each of these individuals (one R and two S). For each control, we recovered the exact same three sequences in both TA cloning and long-read sequencing. This demonstrates the robustness of Namias *et al.’*s (2022) approach, which could thus be used to process much larger numbers of samples in the future.

Overall and across all sampling years, we eventually collected three sequences, one R and two S, for the 28 “triple-peak” individuals from Yopougon and Yammoussoukro. All R copies were identical to D_1_(R) (*i.e.* the R copy carried by D_1_). We also found 26 different S copies, that could be either D(S) or S (see Fig. 1 B “combined PCR. The S copies differed by only a few mutations, resulting in a relatively low sequence divergence (*d* = 0.012), mostly found in introns (*d_exons_*= 0.008 *vs. d_introns_*=0.027).

To try and identify the different D alleles (*i.e.* discriminating the D(S) copies from the S alleles), we compiled a neighbour-joining phylogram with all these S sequences (NB: each “triple-peak” individual carries at least one D allele, and its D(S) and S copies are necessarily different), and combined different informations:

1 We expected D(S) copies to be higher in frequency as they are directly selected for in the presence of insecticides; therefore, clusters of identical S sequences were more likely to correspond to D(S) copies than to S alleles (see Supp. Info. Fig. 2). Moreover, when several individuals had a first S sequence in a cluster, their second S copies would usually be different, and thus attributed to single-copy S alleles.

**Figure 2.**
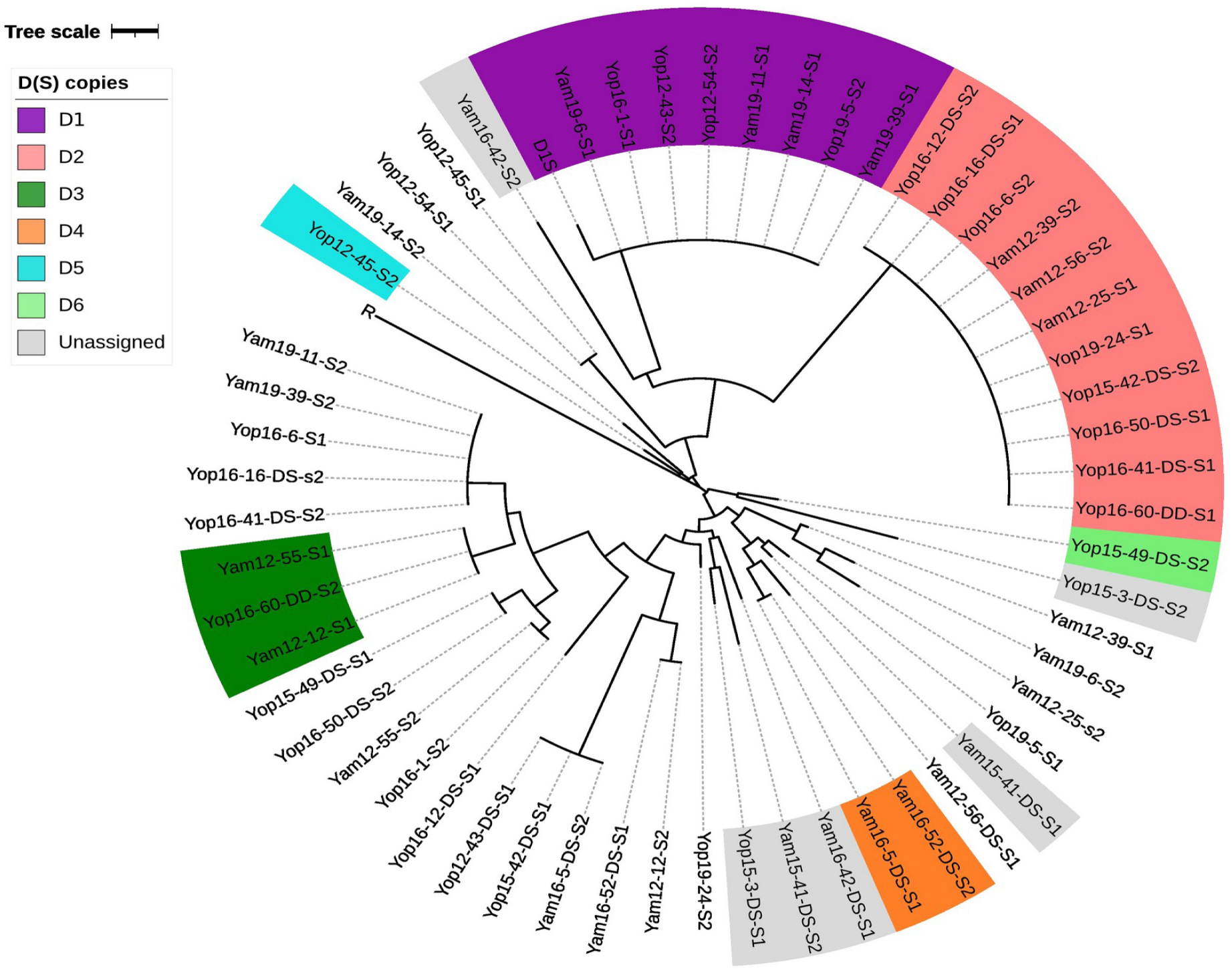
Diversity of the *ace-1* S and D(S) sequences in individuals diplaying triple peaks. This phylogram represents the diversity of the S copies retrieved (TA cloning or Nanopore sequencing) from individuals displaying triple peaks in the mix sequence of the “*ace-1* resistance phenotype” PCR product (see Materials). Samples are coded as follows: locality (Yam for Yamoussoukro and Yop for Yopougon)/sampling year and individual number (-x). 12 samples whose genotype was obtained through short read sequencing are annotated with DS (duplicated heterozygote) or DD (duplicated homozygote). For each individual, the two S copies are indicated as copy S1 and copy S2 (assigned randomly). Sequences indentified as single-copy S alleles are not highlighted. Copies indentified as probable D(S) copies (see text) are highlighted according to the corresponding putative D allele (legend). Unassigned sequences, *i.e.* S sequences that are found in an individual carrying a D allele, but that cannot yet be assigned to S or D(S) for lack of data, are highlighted in grey.

–A first cluster, corresponding to D_1_(S) (purple, Fig. 2), was expected, and found (note that the sequences differed by one mutation from the canonical D_1_(S), including for the D_1_ controls. The existence of this D_1_(S) cluster, and the fact that individuals carrying this D_1_(S) copy carried second S copies that were different, validated our approach. For example, see the individuals Yam19-11 and Yam19-24 (NB: Individuals are coded as follows: locality – Yam for Yamoussoukro and Yop for Yopougon –/sampling year– 12, 15, 16 or 19 for 2012, 2015, 2016 or 2019, resp. – and individual number/sequence code – -x –): their shared copy, Yam19-11-S1 and Yam19-14-S1, is similar to D_1_(S), but their other S copies, Yam19-11-S2 and Yam19-14-S2, are different (Fig. 2); they were thus identified as D_1_S individuals.
–Similarly, a second large cluster was found (pink, Fig. 2), which can unambiguously be assigned to a second D allele, thereafter D_2_.
–Two smaller clusters were found, with two to three individuals sharing one S copy, the other being different (and different from D_1_(S) and D_2_(S)), that we tentatively named D_3_(S) and D_4_(S) (dark green and orange, resp., Fig. 2). Note that, rather than inflating the number of potential D alleles, we chose to conservatively consider individuals carrying one sequence similar to D_1_(S) or D_2_(S) as DS heterozygotes, even if the second S copy was found in another cluster, so that they could actually be D_i_D_j_ heterozygotes (e.g. Yam19-39 and Yam19-11 carry either D_1_(S) or D_2_(S), but their second S copies cluster together; Fig. 2), although for three of those individuals (Yop16-41, Yop16-6, Yop16-16) the analysis of *ace-1* depth of coverage (Supp. Info. Fig. 1) actually confirmed that they were actually DS heterozygotes. Following this principle, for Yop12-45, we identified the D(S) copy as being Yop12-45-S2, thus tentatively named D_5_(S) (light blue, Fig. 2), as Yop12-45-S1 was identical to Yop12-54-S1 while Yop12-54-S2 was identical to D_1_(S) (Fig. 2).

1. For some individuals, the total number of *ace-1* copies they carried was independently known from genomic analyses.

–Yop16-60, carries four copies (as shown by *ace-1* depth of coverage, Supp. Info. Fig. 1), but with only three different sequences, one R and two S. This individual thus carries two different D alleles: Yop16-60-S1 was identical to D_2_(S), while Yop16-60-S2 belonged to the tentative D_3_(S) cluster, this individual is thus a D_2_D_3_ heterozygote (Fig. 2). Note that this also confirms the actual existence of the D_3_ allele.
–Conversely, *ace-1* depth of coverage showed that Yop16-50 carries three *ace-1* copies (Supp. Info. Fig. 1), and is thus a DS heterozygote. As Yop16-50-S1 is identical to D_2_(S), it suggests that Yop16-50S2 is a single-copy S allele. It follows that Yop15-49-S1, which is identical to Yop16-50S2 (Fig. 2), is most probably a single-copy S allele too. Consequently, as Yop16-49 carries a duplicated allele (it carries one R and two S copies), it is its second S copy, Yop15-49S2, that is the D(S) copy (despite being isolated in the tree; Fig. 2); this allele has thus been named D_6_.
2. For the last three D-carriers, Yam15-41, Yam16-42 and Yop15-3, both S copies were singletons in the tree (Fig. 2). Thus, although we know that they each carry at least one new D allele (D_7_, D_8_ and D_9_), which of their two S sequences corresponds to D(S) is not yet determined.

Over our 28 triple-peak individuals, we were thus able to conservatively infer that at least nine different D alleles were segregating in Yamoussoukro and Yopougon (including D_1_, Fig 2 and Supp. Info. Tab. 4), but apart from D_1_, D_2_ and D_3_, the identification of their D(S) sequence remains tentative. Of course, due to limited and non-exhaustive sampling, this is but a minimum estimate of the real diversity of duplicated alleles. Moreover, this diversity of D alleles in Ivory Coast is not recent: we detected D _1_, D_2_ and D_3_ in samples collected in 2012 and repeatedly thereafter (Supp. Info. Tab. 4), suggesting a long-lasting persistence of these alleles. The apparent punctual presence of the remaining D alleles should not be misunderstood for a clue to their fleeting prevalence, as the protocol of our study was designed to assess the overall diversity of D alleles, but not to infer their frequency, or even their presence over time (the sampling size is too limited, 1 to 8 individuals per year and population, and limited by triple peak signal).

This high diversity of *ace-1* D alleles segregating in only two populations from Ivory Coast immediately begs the question of its origin, *i.e.* whether a single original duplication event followed by secondary rearrangements (e.g. recombinations, deletions) might have generated this diversity, or if multiple independent events of duplication, one for each allele, were required. To try and answer this question we used two approaches, the first one geographical, the second genomic.

### *ace-1* sequences do not show structure at the geographical or species level

We first tried to assess whether the different D alleles could be tied to a particular geographical origin in West Africa. We thus added ≈30 *ace-1* S sequences from neighbouring countries (Benin, Burkina Faso, Ghana) of both *A. gambiae* and *A. coluzzii* and computed a phylogenetic model (Fig. 3; Tamura 3 parameters G+I, see Materials). This revealed no clustering pertaining to geographical origin or species of the different S copies, whether S or D(S), despite testing different models of evolution: we found that close, or even identical, S copies can be found in all countries, and that none of the different D(S) copies is tied to a particular country.

**Figure 3.**
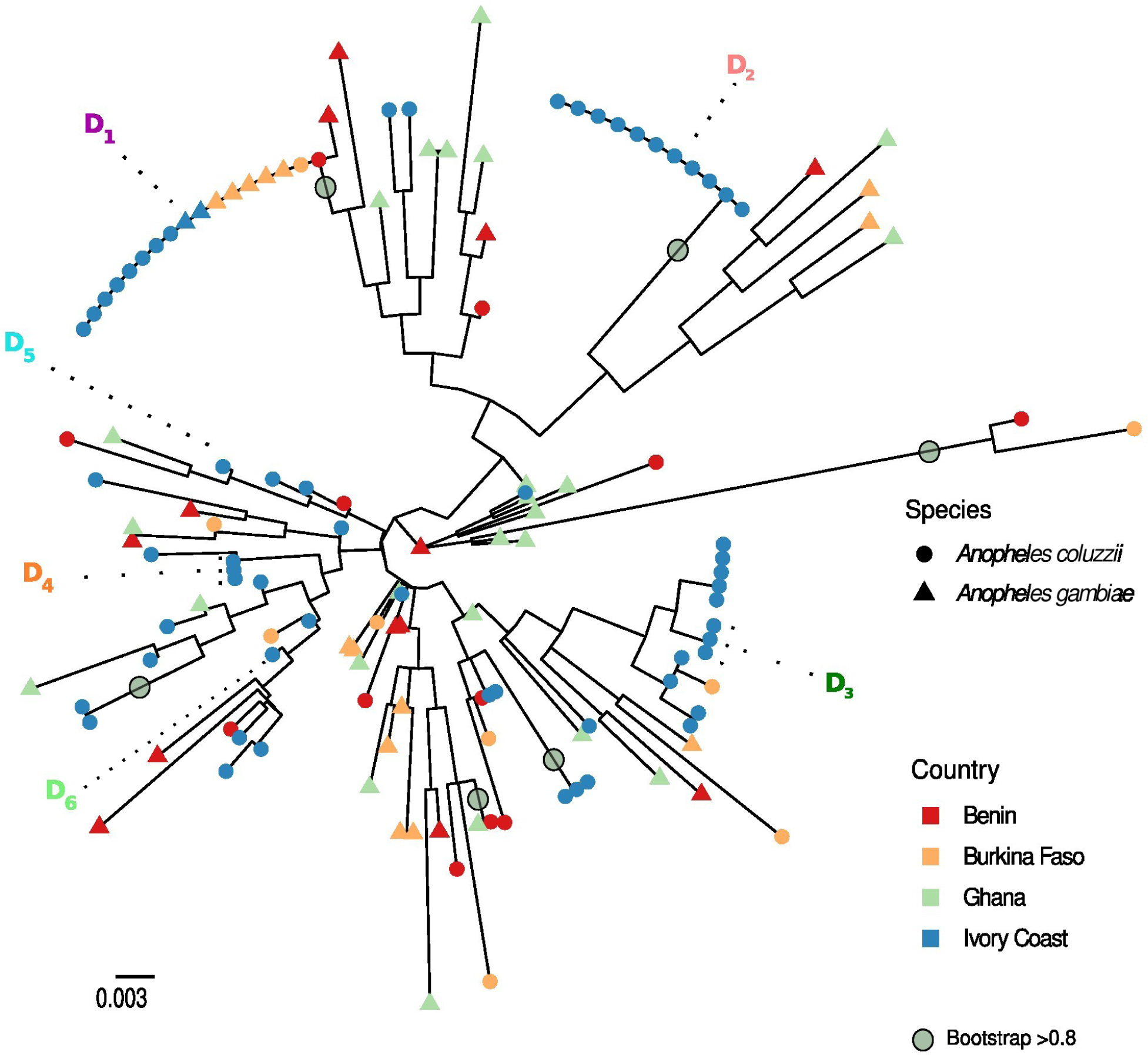
*ace-1* S diversity in West Africa. Maximum likelihood tree of *A. coluzzii* and *A. gambiae* S sequences from West Africa. For each sequence, we indicated its species (triangle for *A. gambiae* and circle for *A. coluzzii)*, its geographical origin (Benin in red, Burkina Faso in orange, Ghana in green, and Ivory Coast in blue, as in the inlet map); for references see Supp. Info. Tab. 2. The D(S) copies indentified in the present study are also indicated, as well as the bootstraps ≥0.8 whith gray circles.

In fact, the only highly supported nodes are those of the D(S) clusters (Fig. 3). The diversity is indeed relatively low and mostly concentrated in the introns. Translating the exonic part of the *ace-1* sequences showed that almost all mutations were synonymous. The only notable exceptions were found in three samples from Ghana (not shown), although the impact of those mutations is entirely unknown.

We have thus so far no evidence as to whether the different D alleles identified in the present study have geographically originated in the very populations where we found them or elsewhere, and we cannot rule out the possibility of a single origin followed by secondary recombinations.

### All D alleles share a common genomic architecture

As the geographical approach failed to reveal the origin of the D alleles, we took advantage of the bioinformatic approach developed by Assogba *et al*. (2016) to address this question by analysing the genomic architecture of these alleles: we already knew that D_1_ and R^x^ alleles share the same breaking points, but what about these new alleles? Different genomic structures, *i.e.* different amplicon sizes and breaking points, for the D alleles would indeed support independent origins.

Illumina-generated 350 bp-spaced paired-end reads (150 bp) of twelve individuals carrying different D alleles (D_1_, D_2_, D_3_, D_4_, D_7_, D_8_, D_9_. we could not get reliable DNa extraction for sequencing individuals carrying D_5_ and D_6_) were first mapped onto the reference *A. gambiae* PEST genome (Vector-Base; AgamP4.13); we also mapped the Kisumu genome as a non-duplicated reference. For each individual, using the mean depth of coverage (*µ*DoC) over the full chromosome as reference, we then calculated the standardized depths of coverage (*pb*DoC*_std_*) of each base between positions 3,436,000 and 3,639,000 of the 2R chromosome (*ace-1* lies between positions 3,484,107 and 3,495,790; see Materials and Methods). As expected, Kisumu’s *pb*DoC*_std_* remained close to 1 over the whole region (Fig. 4A). By contrast, all D-carriers displayed a consistent *pb*sDoC*_std_* increase over a 203 kb region encompassing the *ace-1* locus (between positions 3,436,000 and 3,639,000), very similar to that observed for D_1_ (Fig. 4B and Supp. Info. Fig. 1). We observed for 11 individuals a 1.5-fold *pb*sDoC*_std_* increase (1.5 ±0.14 for the duplicated region *vs.* 1 ±0.16 for the flanking non-duplicated regions), which is consistent with a DS genotype (Supp. Info. Fig. 1). Yop16-60, displayed a 2-fold *pb*sDoC*_std_* increase in the same area (Fig. 4B), suggesting a DD genotype (sequences revealed the D_2_D_3_ genotype, see D(S) copies identification above).

**Figure 4.**
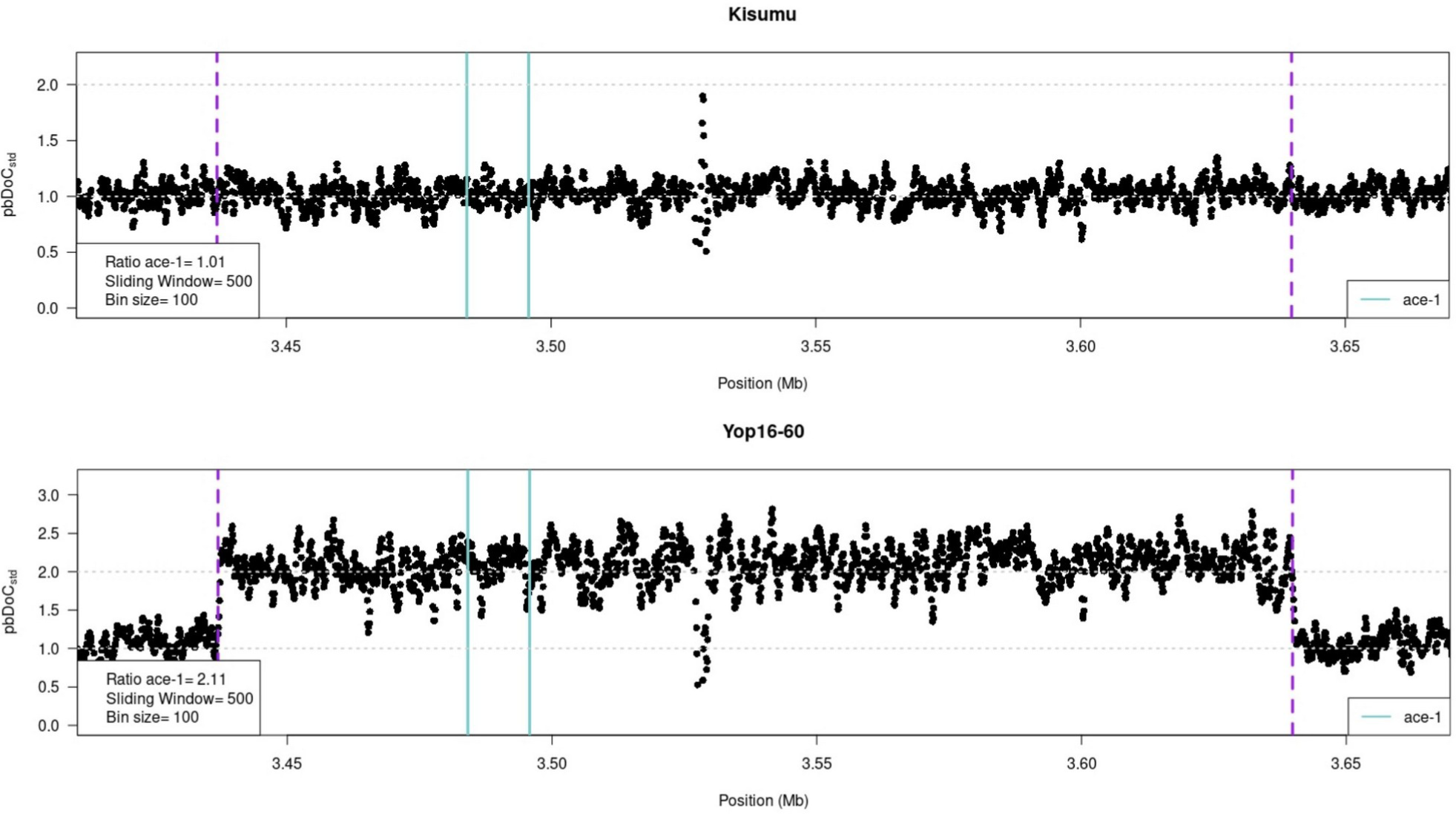
Genomic architecture of the *ace-1* D alleles. In each graph, we presented the variation of the standardized per-base depth of coverage (*pb*DoC*_std_*, with 1 being the mean *pb*DoC calculated over the whole chromosome) along the chromosomal region of interest (absiss, from 3.4 to 3.7 MB along the chromosome 2R). Each dot is the mean *pb*DoC*_std_* calculated every 100 bases (bin size) over 500-base sliding windows. The purple dashed lines represent the amplicon limits of the D_1_ and R^x^ alleles (Assogba et al. 2018); the cyan lines represent the *ace-1* gene location. A-The susceptible strain Kisumu (upper graph) is the single-copy S allele reference, with no particular variation of *pb*DoC*_std_* (mean = 1). B-The lower graph represents the *pb*DoC*_std_* variation for the individual Yop16-60 (as a representative example of the D alleles analyzed in the present study; similar graphs for the other individuals analyzed are shown in Supp. Info. Fig. 1). A 2-fold increase reveals the amplicon size and location, similar to D_1_; it is consistent with a D_i_/D_j_ genotype (two S and two R copies). All D_i_ alleles share the same breakpoints as D_1_; however, the other individuals display only a 1.5 increase, as expected for genotypes DS (two S and 1R copies; Supp. info. Fig. 1).

To determine the precise position of the duplication breakpoints, we isolated reads mapping at ±1 kb from the putative breakpoints determined from the DoC graphs. All individuals showed a significant increase of soft-clipped reads, *i.e.* reads overlapping the 203 kb-amplicon junction, on positions close to the 5’ and 3’ breakpoint positions previously identified for D_1_ (position 3,436,927 and position 3,639,836; resp.; Fig. 4B; Assogba *et al*. 2016), further suggesting very close, if not identical, breakpoints for all the analysed D alleles (D_1_, D_2_, D_3_, D_4_, D_7_, D_8_, D_9_). The size of the duplication was further confirmed to 203 kb using the distribution of the insert-size of the reads overlapping the region.

We finally submitted these individuals to the specific PCR test designed by Assogba et al. (2016) that amplifies a 460 pb segment overlapping the junction between the amplicons (similar between D_1_ and R^x^ in Assogba et al. 2016). The segment fragment was amplified in all 12 individuals, further confirming that the 3’ breakpoint location is conserved for the various D alleles tested.

Together, this PCR test and the DoC analysis indicates that the amplicons for all the D alleles analysed (D_1_, D_2_, D_3_, D_4_, D_7_, D_8_, D_9_) share exactly the same genomic architecture, *i.e.* the same boundaries and the same sizes, without internal deletions (as seen in R^x^ alleles, Assogba et al. 2018).

## Discussion

We investigated a significant departure from expected heterozygosity at the *ace-1* locus in two populations of *A. coluzzii* of Yamoussoukro and Yopougon (Ivory Coast). As this excess could not be entirely explained by the presence of the only known heterogeneous duplicated allele, we went through a genotyping and sequencing (short and long reads sequencing and TA cloning) protocol to sort out putative new *ace-1* duplicated alleles. We found at least eight new D alleles that shared the same structure and R copy, but carried different S copies; moreover, it seems that several D alleles have been segregating in these populations since at least 2012.

### An unsuspected duplicated allele diversity

First thought to be limited to a unique heterogeneous duplication (D_1_) and multiple homogeneous ones (R^x^) across all West Africa (Assogba *et al*. 2016), we now have solid proof that the diversity found at the *ace-1* gene is much larger. The existence of multiple R^x^ alleles is expected as unequal recombination is frequent for multiple-copy genes (e.g. esterases, Milesi *et al*. 2016). Their persistence in the populations also can be easily explained, as these alleles have been proved to have different fitness impacts directly linked to their copy number, *i.e.* a greater number of R copies results in higher resistance levels, but also to stronger selective disadvantages in the absence of insecticide (Assogba *et al*. 2016). A large diversity of R^x^ copy-number-alleles thus reflects as many trade-offs with different levels of resistance and a wide array of R^x^ duplications can be maintained in natural populations exposed to high but variable insecticide treatment intensity. This however, cannot be said of D alleles: they confer a heterozygote phenotype, and can be selected in areas where insecticide treatments are not always present (either temporally or spatially); however, they are only variable in the sequence of the S copy, which is not expected to modify the original trade-off, as indicated by previous studies of such alleles in *C. pipiens* (Labbé et al 2007, Milesi et al 2018).

This analogy with *C. pipiens* could however provide an explanation for the persistence of D alleles diversity. In *C. pipiens* mosquitoes, many populations are polymorphic for D alleles. These alleles are sublethal when homozygous (D_i_D_i_), probably due to different mutations embarked in the duplication, as they complement when heterozygotes (D_i_D_j_) (Labbé *et al*. 2007 and Milesi *et al*. 2018). Milesi et al. (2018) have shown through modeling that different sublethal D alleles that complement each other can be maintained in the same population by frequency-dependent selection: as they are only detrimental when homozygotes, but initially present mostly in heterozygotes, they are first selected, increase in frequency, but rapidly reach a stable plateau when frequent enough to generate homozygotes. Note that this equilibrium can only be reached with sublethal D alleles, the introduction of a single non-sublethal D allele leading to its fixation (Milesi et al. 2018). In the case of *A. gambiae s.l.*, *Ag*-D_1_ was not found sublethal when homozygous in lab experiments (Assogba et al. 2015), so one may wonder if the same mechanism as the one proposed for *Culex* is at work. Moreover, the overall D frequency remained surprisingly stable from 2012 to 2019, despite a slight increase in resistance (binomial test, *p =* 5.5×10e^-11^ and *p* = 2.4×10e^-4^ for Yopougon and Yamoussoukro, respectively; Supp. Info. Fig. 2). The fitness impact of the new D alleles thus remains to be investigated, and only through its characterization could we explain the persistence of their diversity.

### Beyond the spotlight effect

Since Djogbénou *et al*. (2008) seminal work, D_1_ had been the only D allele described in *A. gambiae*, despite regular surveys (Djogbenou *et al*. 2008; Assogba *et al*. 2016, 2018; Grau-Bové 2021, Kouamé *et al*. 2023). This highlights a bias in the way these variants are studied: the “classical” approach is based on field-caught individuals crossed in the lab with a reference susceptible strain (Labbé *et al. 2007*; Assogba *et al*. 2016; Milesi *et al*. 2018), will only catch individuals frequent enough to be sampled, and more importantly fit enough to survive and reproduce; this is potentially a major problem when studying duplicated alleles that are often plagued with strong deleterious effects (Innan & Kondrashov, 2010; Schrider & Hahn, 2010; Schrider *et al*. 2013). On the other hand, surveys based on specific molecular tests based on a few diagnostic mutations, are prone to a strong “spotlight effect”, *i.e.* they can only find what they are looking for, especially when these mutations are not directly causal for the resistance phenotype but rather neutral variations.

Because of their inherent ease of data production and their fast application to large sample size, bioinformatics analyses based on NGS data are a promising tool for more exhaustive surveys of natural populations. However, they also come with limitations that should be accounted for: sequencing technologies have come on in bounds and leaps in the last years, improving the quality and length of reads they produce, but they are still prone to errors that may greatly affect studies where certainty and precision is required to genotype wild-caught samples, as is the case in our study. For example, Grau-Bové et al. (2021) suggested variation in the numbers of R and S copies (*i.e.* copy-number-alleles), in Ivory Coast, that we found no evidence of, as all the D alleles identified were carrying only one S and one R copy. This discrepancy probably comes from the fact that, when the identification of S to R copies ratio is dependent on a single position coverage (e.g. the G280S point mutation, see Grau-Bové *et al*. 2021), DOC variations may quite often lead to inaccurate genotyping and copy-number estimations. Similarly, despite indications that suggested the potential for D sequence-alleles, Grau-Bové et al.’s study did not allow their specific identification, in particular because SNP phasing from linkage disequilibrium is difficult with short-reads data, and even more so when several copies are present. When a precise determination of haplotypes spanning hundreds of base-pair of the different copies is required, short-reads are indeed clearly not the best option. We demonstrated the potential of long-reads sequencing to overcome this issue, and as long-reads sequences become more accessible and reliable, especially for SV detection (Mantere *et al*. 2019; De Coster *et al*. 2023, Namias *et al*. 2022), some of these limitations may soon be lifted.

### The origins of D alleles remain to be confirmed

The striking D allele diversity found in only two populations raises questions on the origin(s) of the duplicated *ace-1* alleles in *Anopheles* mosquitoes (Fig. 5). Their strict structural homology, as all share the exact same boundaries and breakpoints, suggests that a surprisingly frequent and localized process of *de novo* unequal crossing-overs in heterozygous individuals would be required to back-up a multiple-event origin (scenario 1, Fig. 5). While Assogba *et al*. (2016) did find a *harbinger* transposable element on the far 3’ end of the duplication that might explain these recurrent recombination events in the same genomic area, the high number of D alleles, together with the R^x^ ones, make this scenario far from parsimonious. The alternative scenario (scenario 2, Fig. 6) indeed requires only one unique first unequal crossing over event, followed by secondary recombination events between the D(S) copy bound in the duplication and a single-copy S allele in heterozygous DS individual. These secondary recombination events could be limited to the *ace-1* sequence or much larger (up to 203kb, *i.e.* encompassing all embarked genes of the duplication).

**Figure 5.**
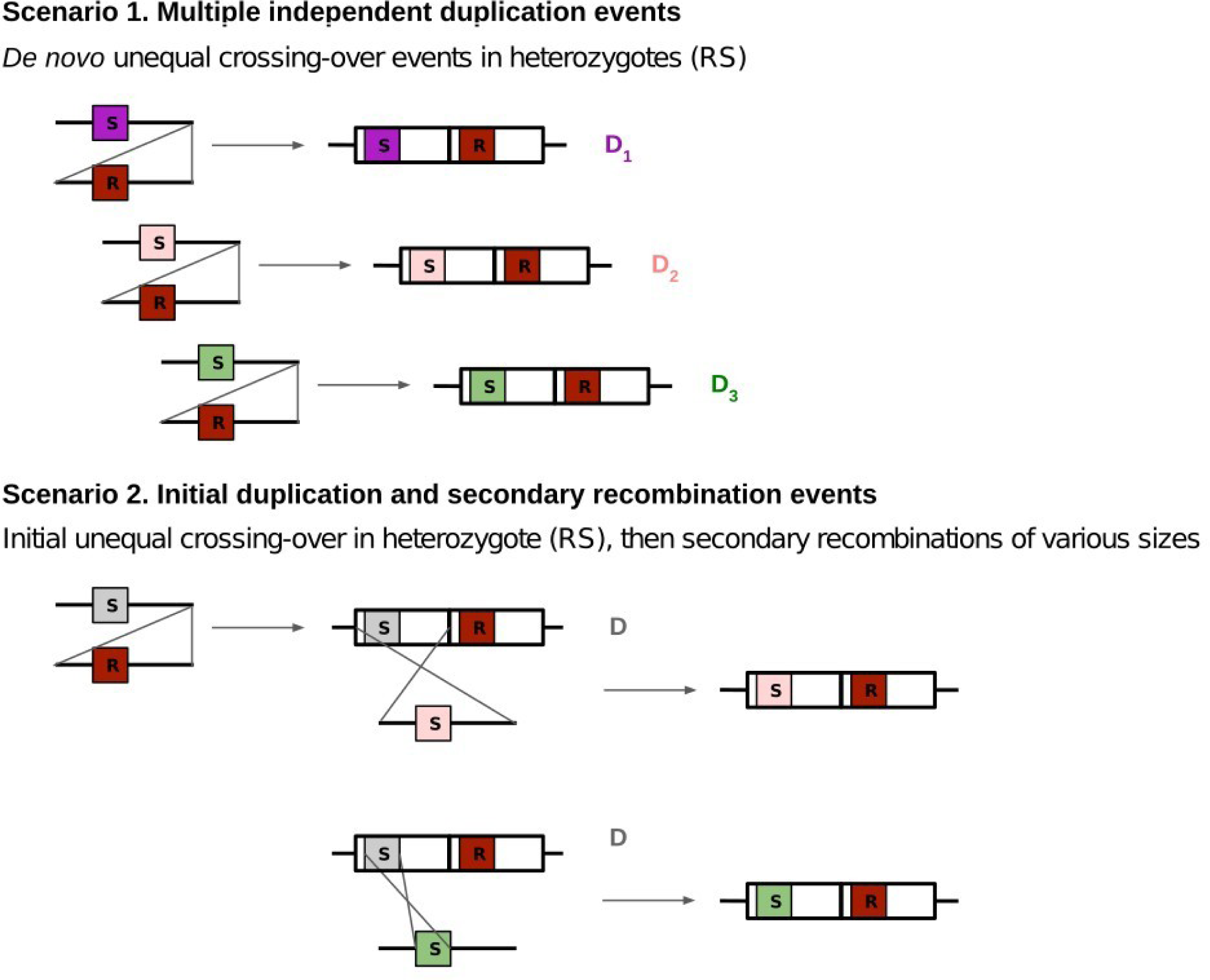
Possible scenarios for the origin of Ivory Coast *ace-1* duplications. Scenario 1 requires several independent duplication events on the same breakpoints, whereas scenario 2 considers a first duplication event followed by secondary recombinations occurring in the amplicon that bears the S copy (either the whole amplicon, as represented for D_2_, or only the *ace-1* locus, as represented for D_3_, or any size between). *NB: the alleles presented here are for illustration only, as the present study did not allow firmly distinguishing the two scenarios, or any secondary recombination span*.

To confidently discriminate those hypotheses however, complete haplotypes of the duplicated alleles are required. Short reads do not allow for a precise phasing of variants called in the duplicated region, and the effective polyploid state of duplicated alleles makes statistical phasing hazardous. However, it should soon be possible, with the improvement of long-read sequencing, to sequence the entire duplicated region. Here we have used an intermediate approach between these two methods, by focussing our efforts on the *ace-1* PCR product sequence, too short to discriminate between the two scenarii, but very efficient to reveal the huge local diversity of duplicated sequence-alleles.

## Conclusion

Altogether our findings highlight the high frequency and diversity of duplicated resistance alleles of the *ace-1* locus in *Anopheles* mosquitoes, probably because of secondary recombination events between single-copy and duplicated alleles, and the need for fitness studies to better understand the mechanisms allowing their persistence in natural populations. However, it also highlights the real challenges to overcome when analyzing such large-scale structural variants (SV). The last 15 years have indeed seen a spur of interest in the role of SVs in the adaptation process (Wellenreuther et al. 2019), thanks to the development of bioinformatics tools and new generation sequencing progress. However, copy-number variations are a perfect example of genomic mutations that are particularly arduous to investigate, due to their inherent genomic characteristics, *i.e.* they are mostly just more of the same sequences, all mapping together on reference genomes. They are thus prone to misinterpretations with too naive approaches. However, these large genomic mutations are frequent and ubiquitous (Emerson et al. 2008; Itsara et al. 2009; Katju & Bergthorsson, 2013; Langley et al. 2012; Reams et al. 2010; Schrider et al. 2013), they played a decisive role in the evolution of living organisms, and are still determinant in the adaptation process, even at the micro-evolutionary scale (Lynch et al. 2008, Katju & Bergthorsson, 2013; Schrider et al. 2013; Kondrashov, 2012); they are thus worth the painstaking endeavour to study them.

---

**Supporting information Figure 1.**
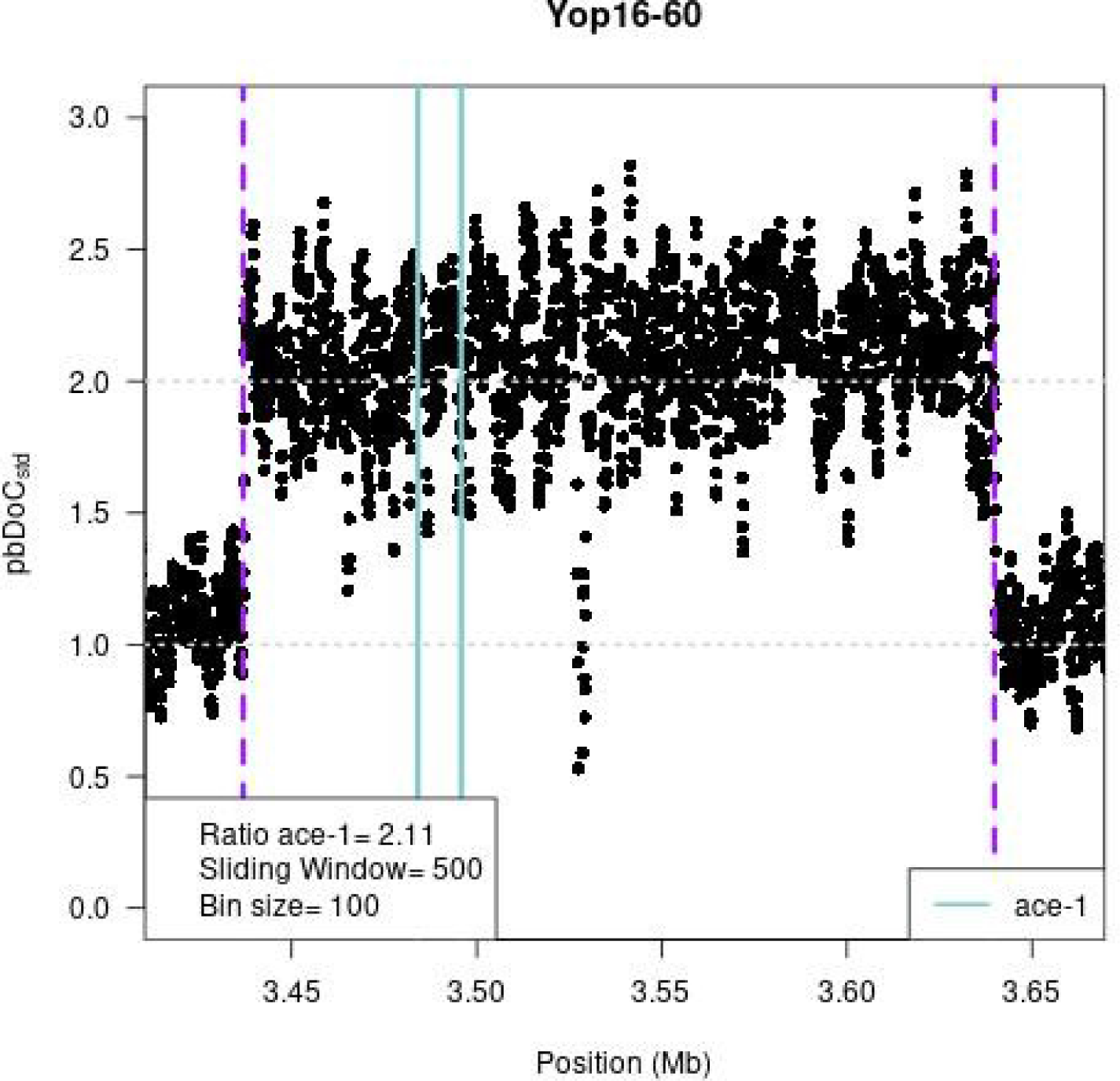
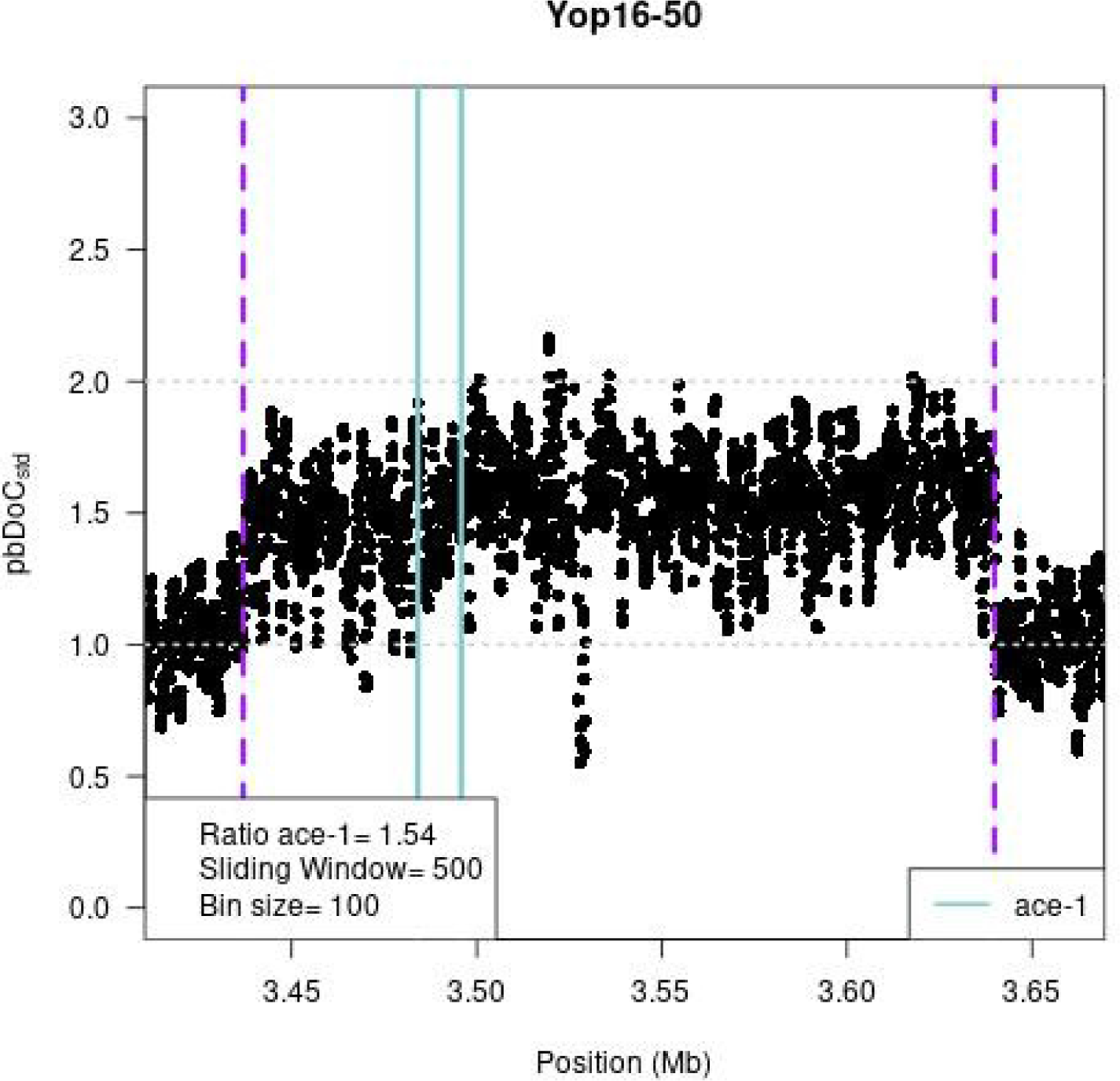
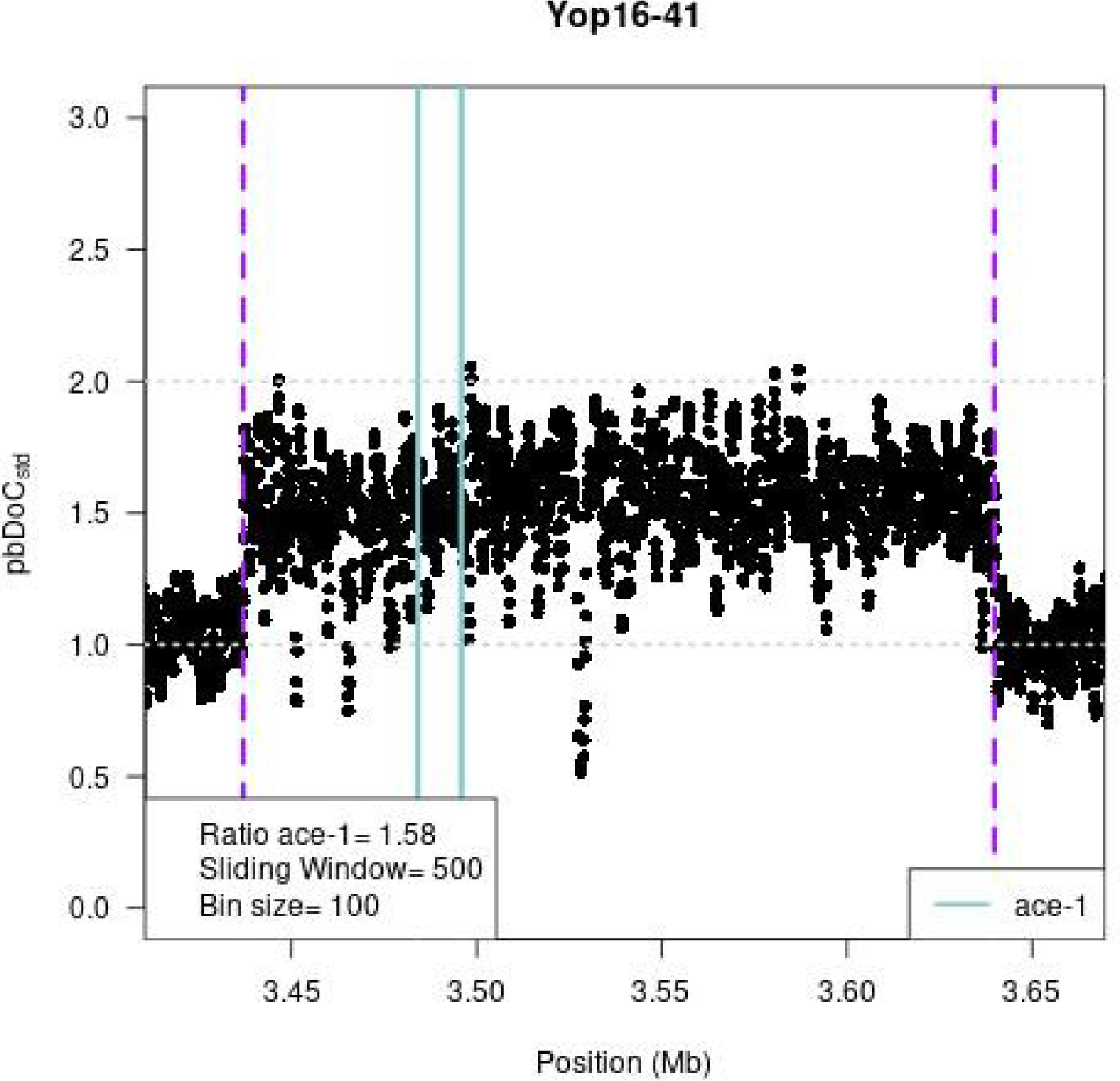
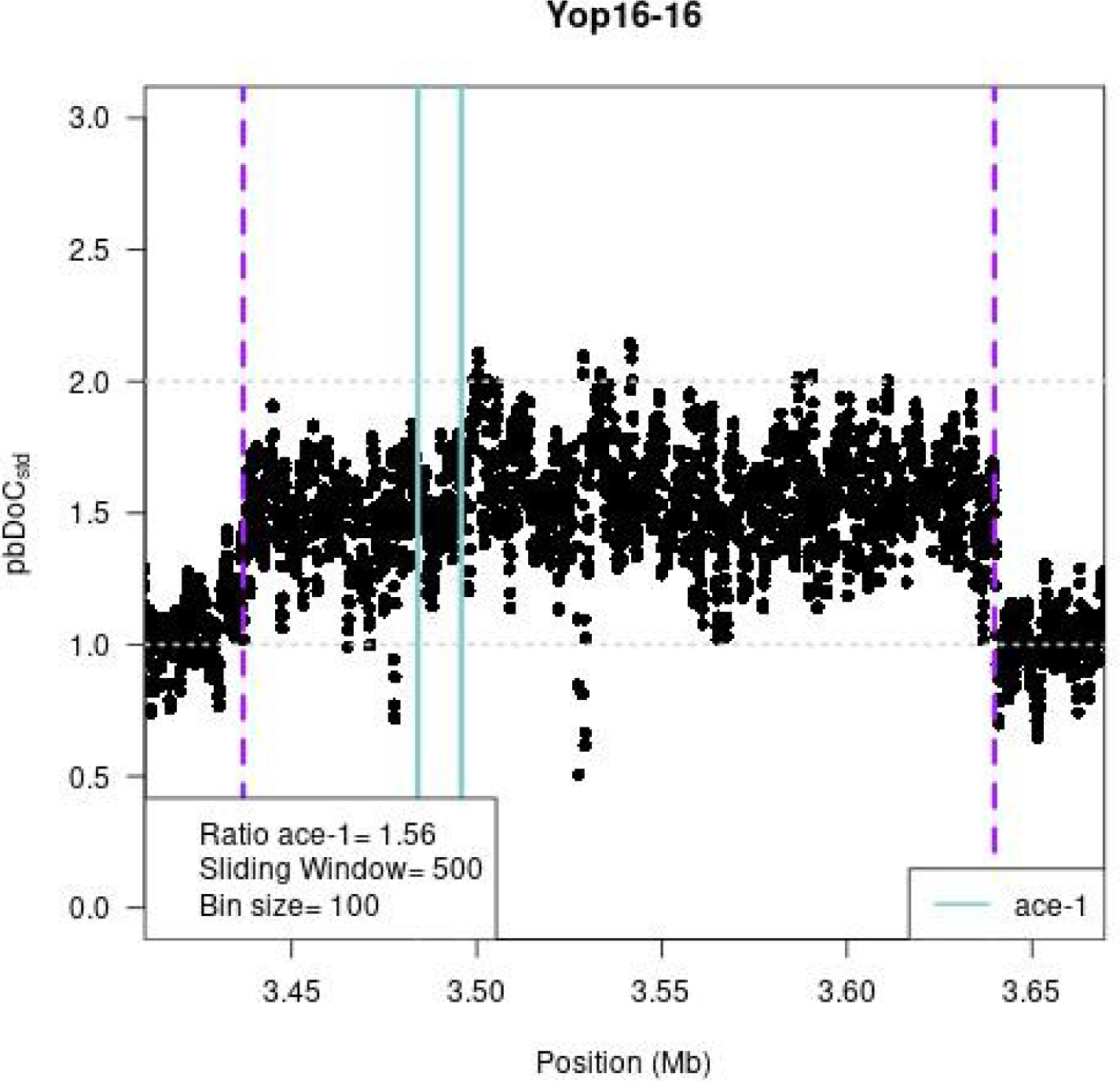
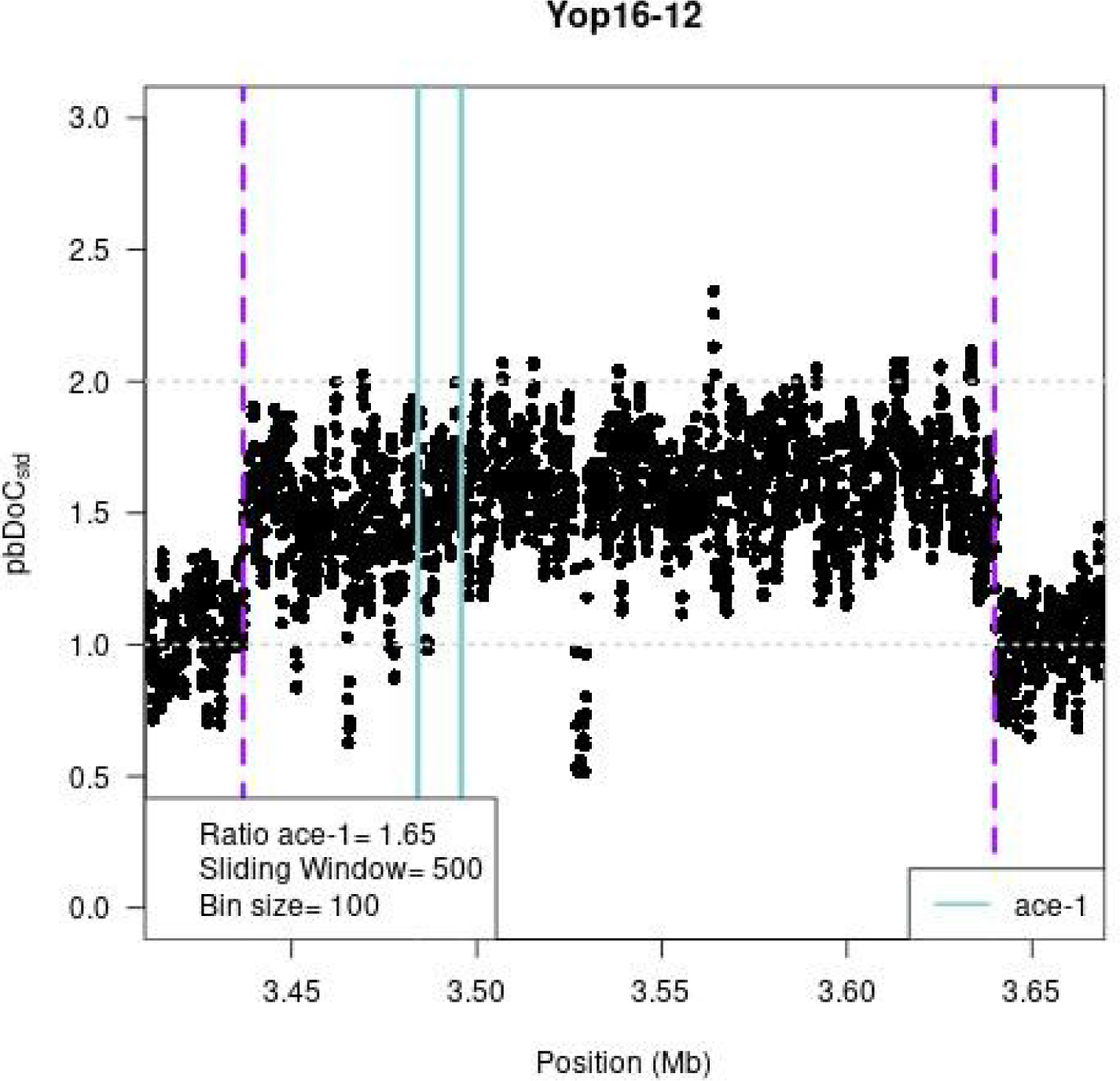
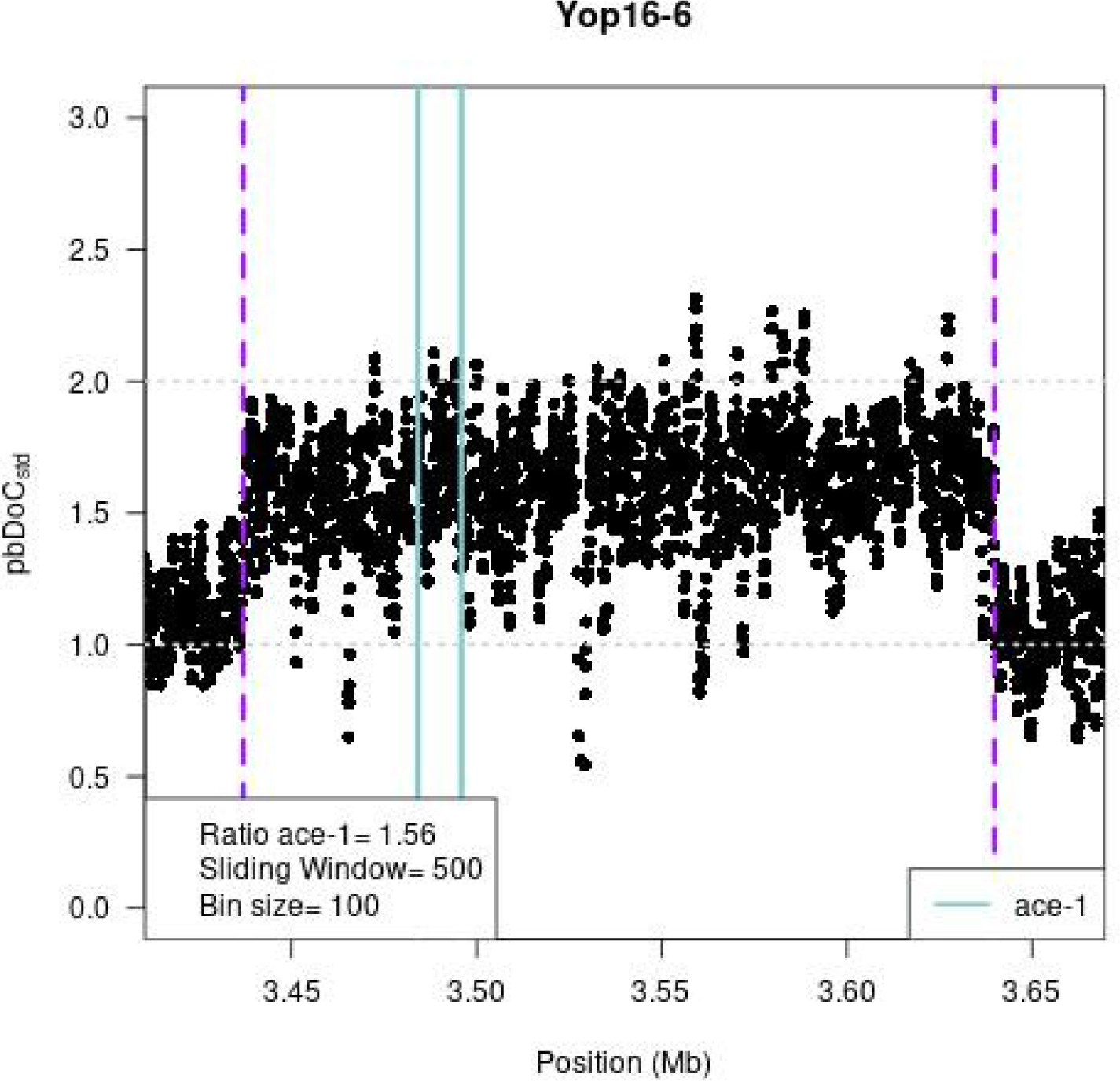
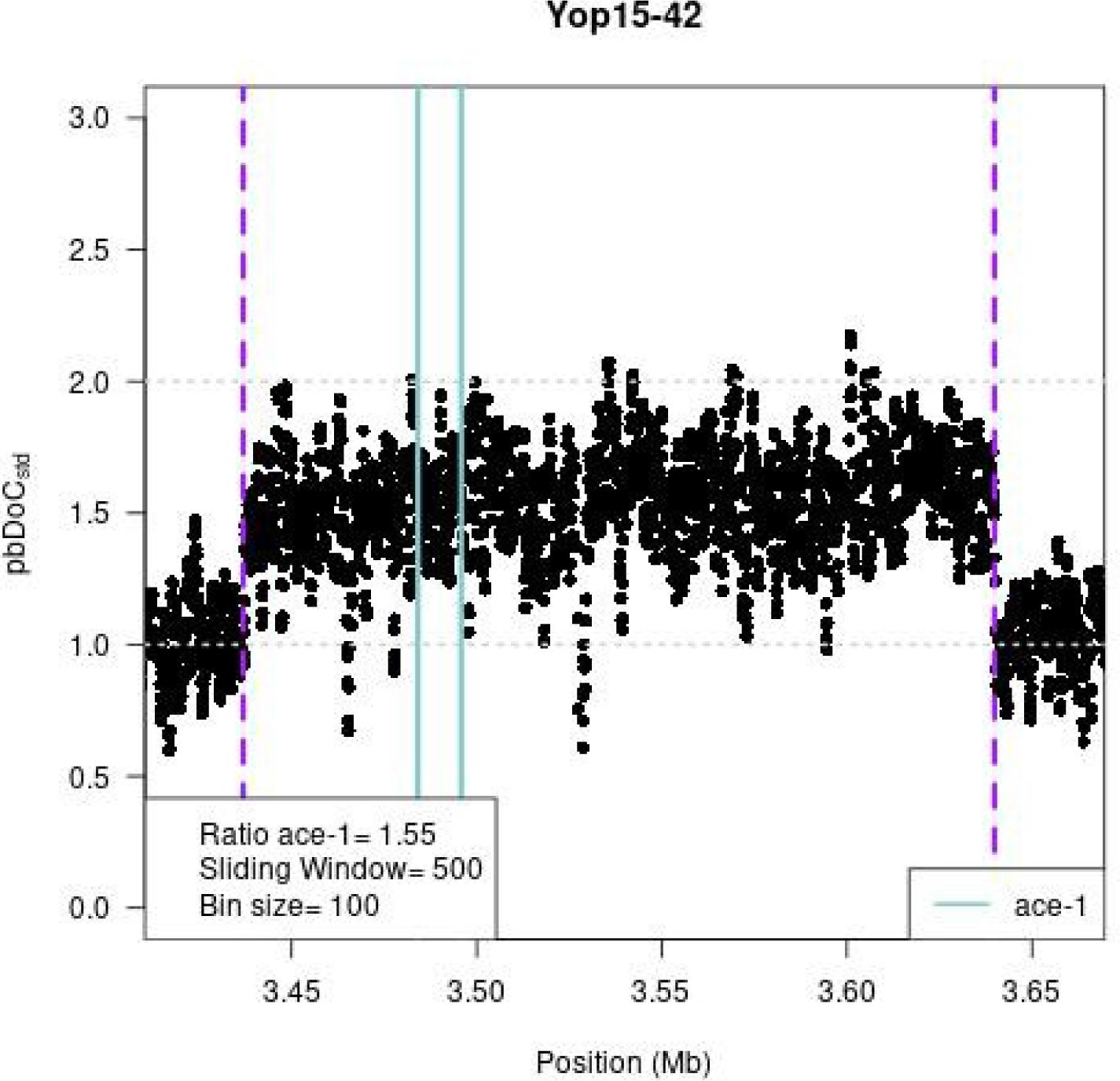
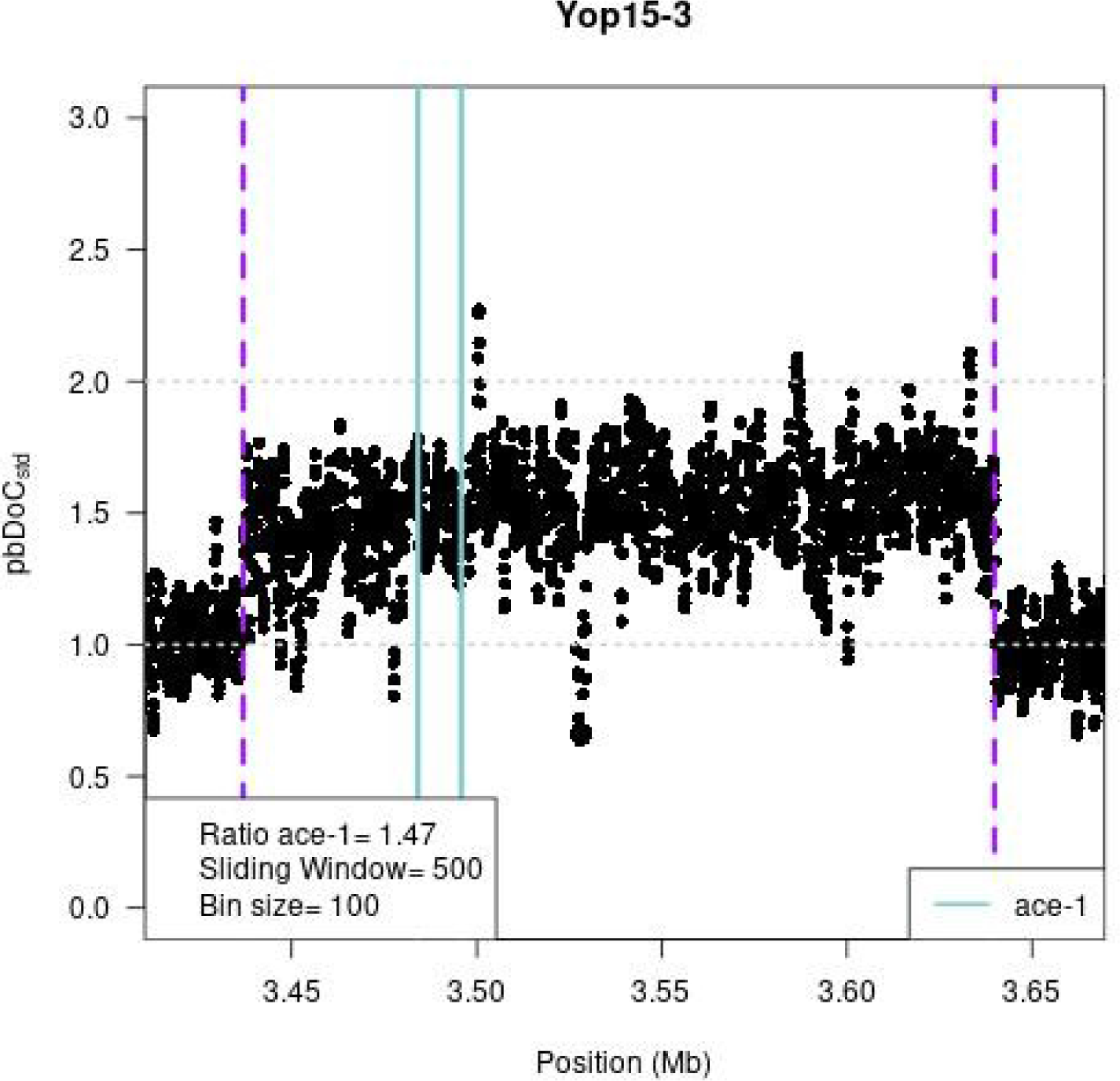
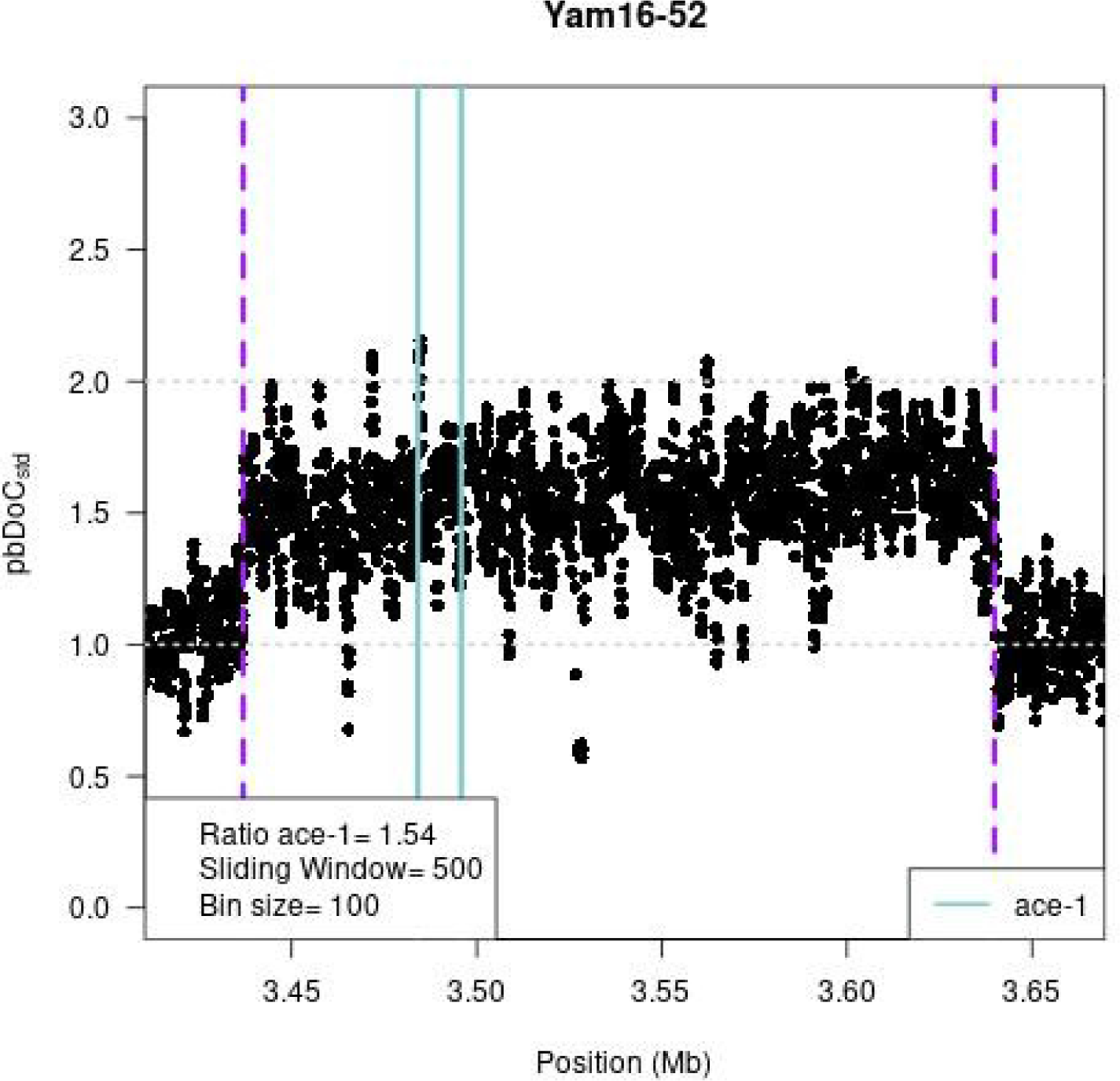
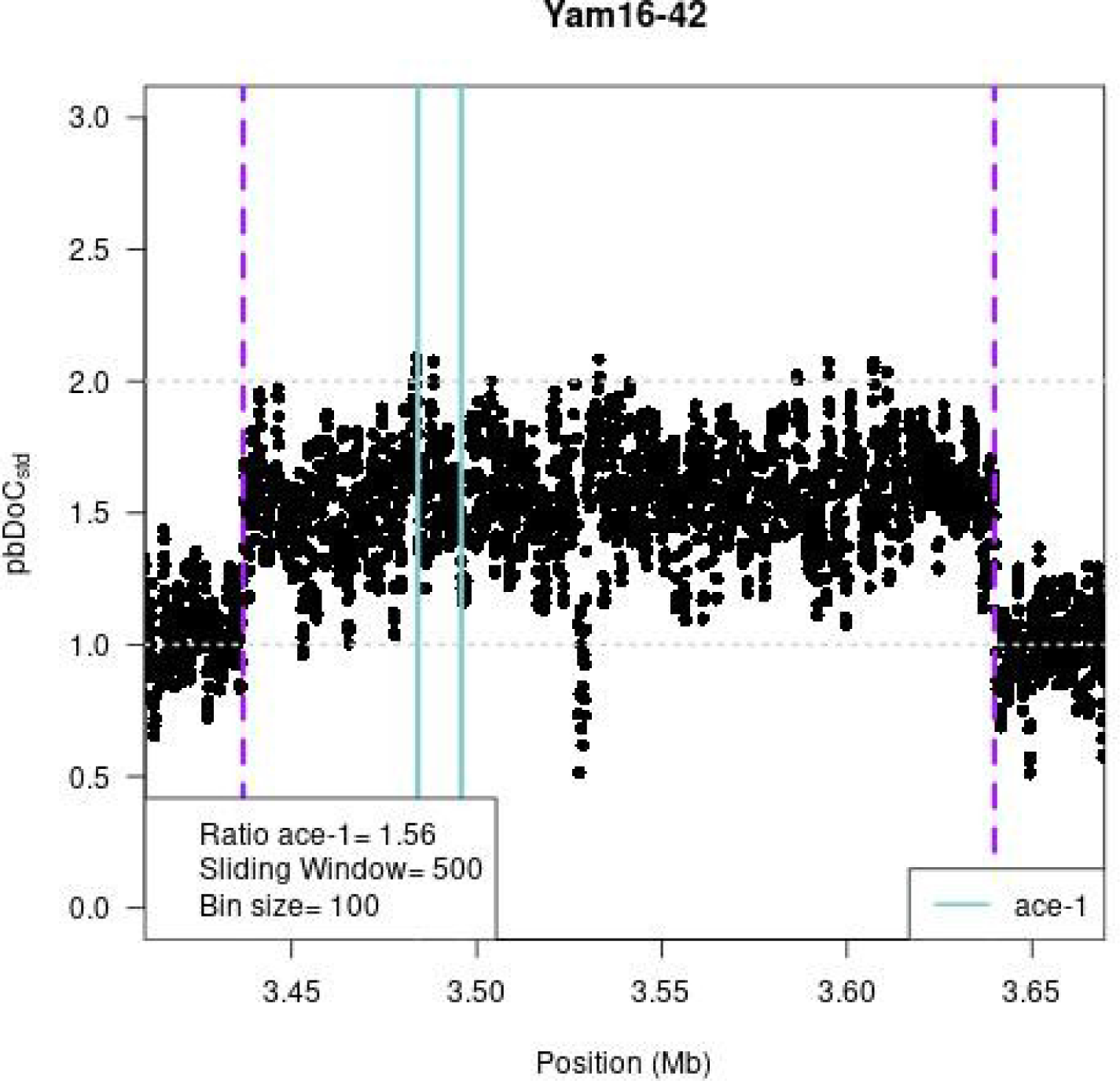
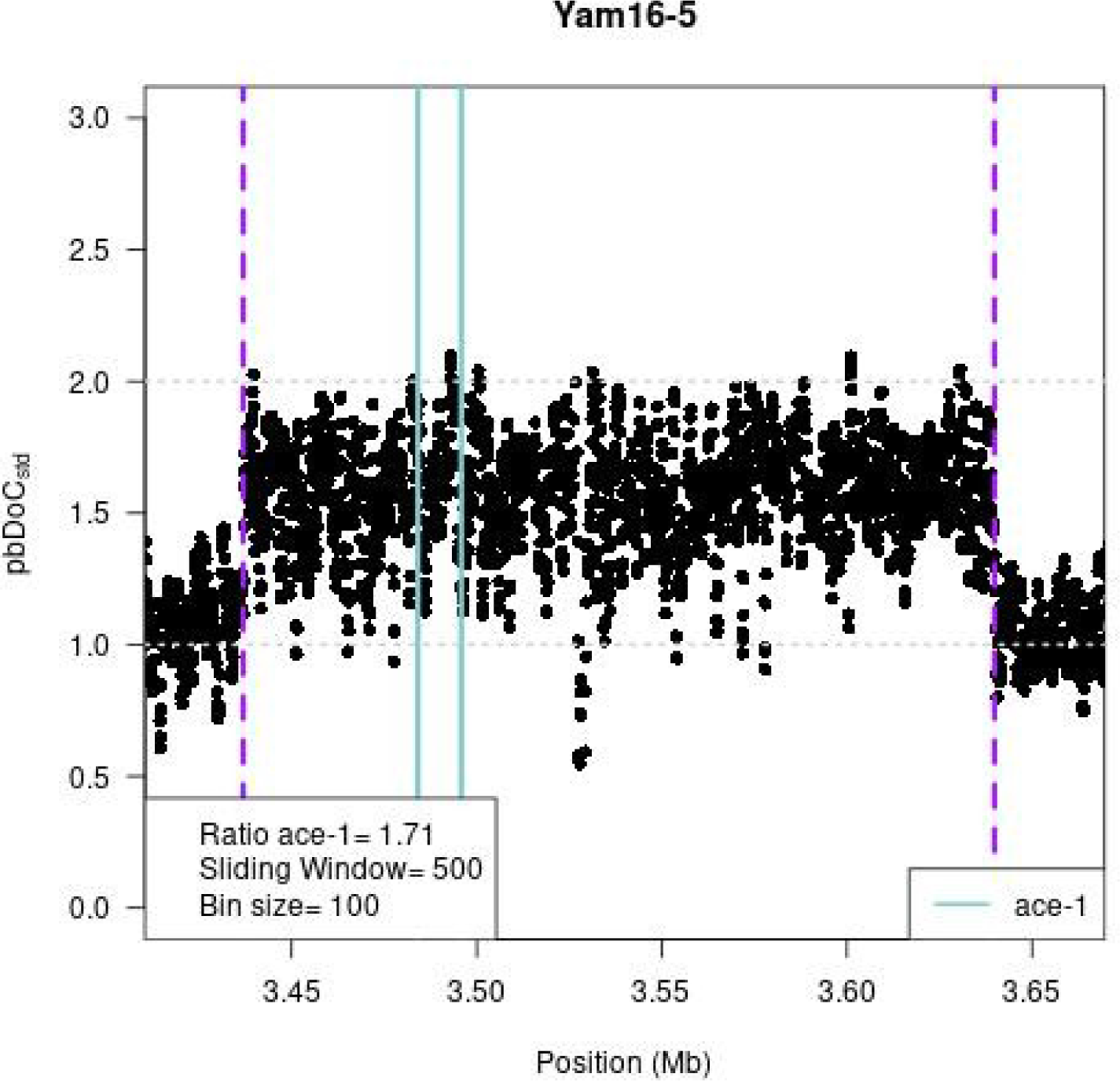
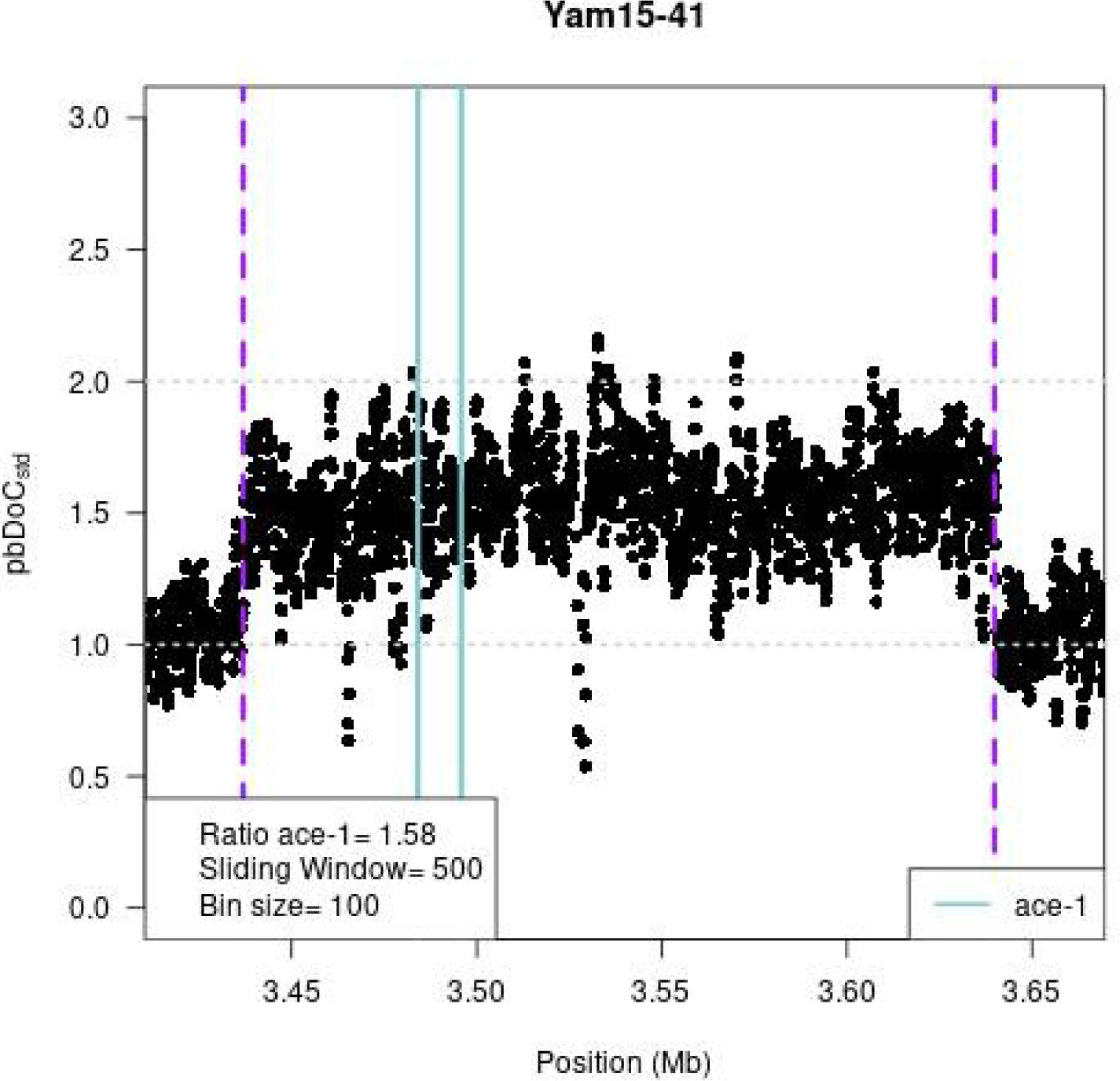
Heterogeneous allele structure. In each graph, we presented the variation of the standardized per-base depth of coverage (*pb*DoC*_std_*, with 1 being the mean *pb*DoC calculated over the whole chromosome) along the chromosomal region of interest (absiss, from 3.4 to 3.7 MB along the chromosome 2R). Each dot is the mean *pb*DoC*_std_* calculated every 100 bases (bin size) over 500-base sliding windows. The purple dashed lines represent the amplicon limits of the D_1_ and R^x^ alleles (Assogba et al. 2018); the cyan lines represent the *ace-1* gene location.

**Supporting information Figure 2.**
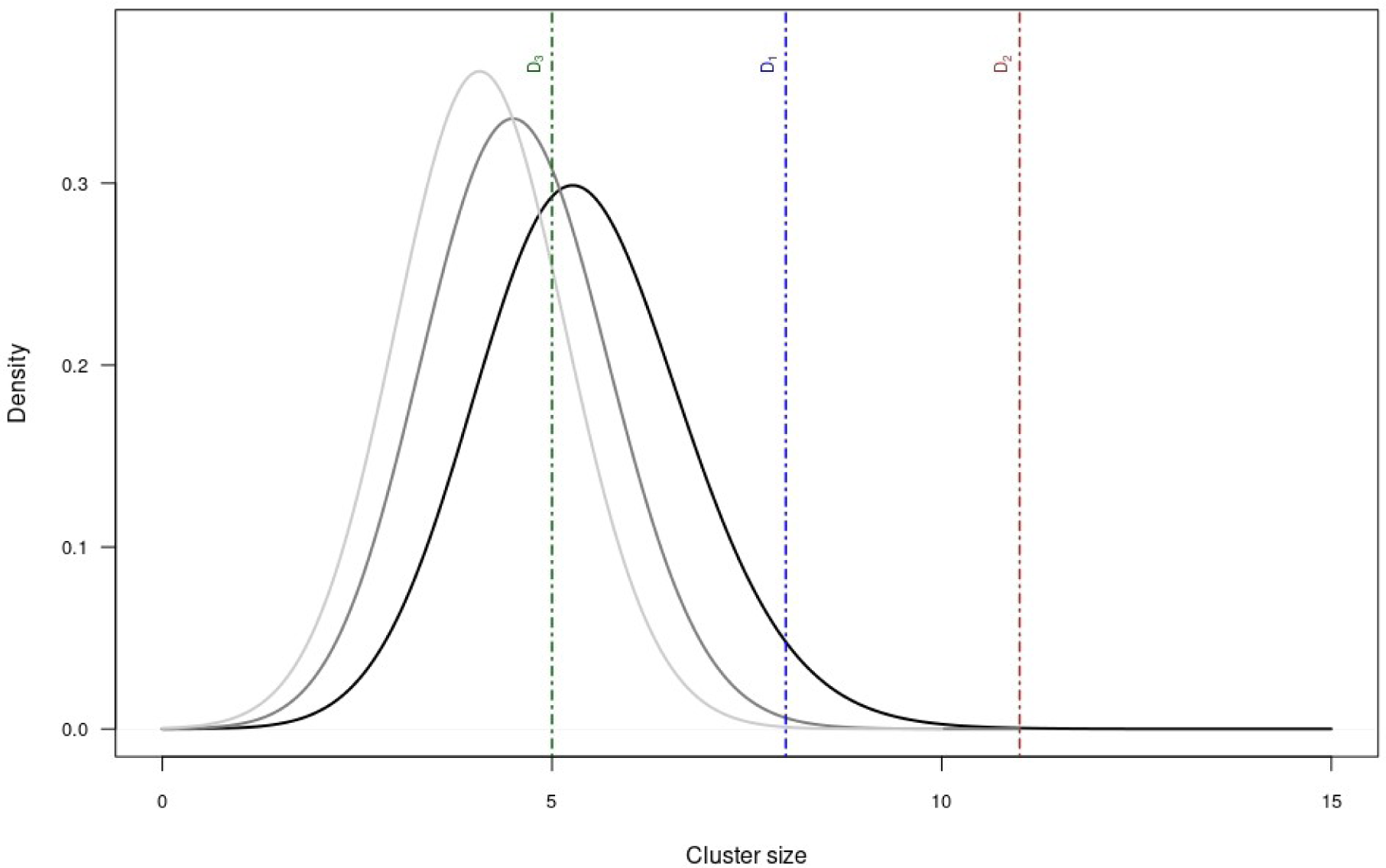
Distribution of expected number of identical sequences in a random draw of 56 sequences (28 2n individuals) out of 26 different S_i_. To test our assumption than D(S) sequences could be recognied because they would be part of larger cluster than single-copy S alleles, we computed the distribution of the expected number of identical sequences in a random draw of 56 sequences (2 per 28 diploïd individuals) out of the 26 different S sequences that composed our dataset, over 100,000 iterations. The probability of observing a given number of sequences in a cluster (*n_obs_*) is given by 1-(the corresponding quantile in the simulated distribution). Solid lines represent the expected number of identical sequences for the first, second and third largest clusters over 100,000 iterations (see text). The dashed-dotted lines are the observed cluster sizes for D_1_, D_2_ and D_3_ (blue, brown and green respectively). Both D_1_ (*n*_obs_ = 9, *p* < 0.001) and D2 (*n*_obs_ = 11, *p* < 0.001) clusters were significantly larger than expected, whereas three-sequence clusters are expected in random draws (*p* = 0.92). Thus, apart from the independently-confirmed D_3_ allele (see text), D_4_(S), D_5_(S) and D_6_(S) identification remains tentative.

**Supporting information Figure 3.**
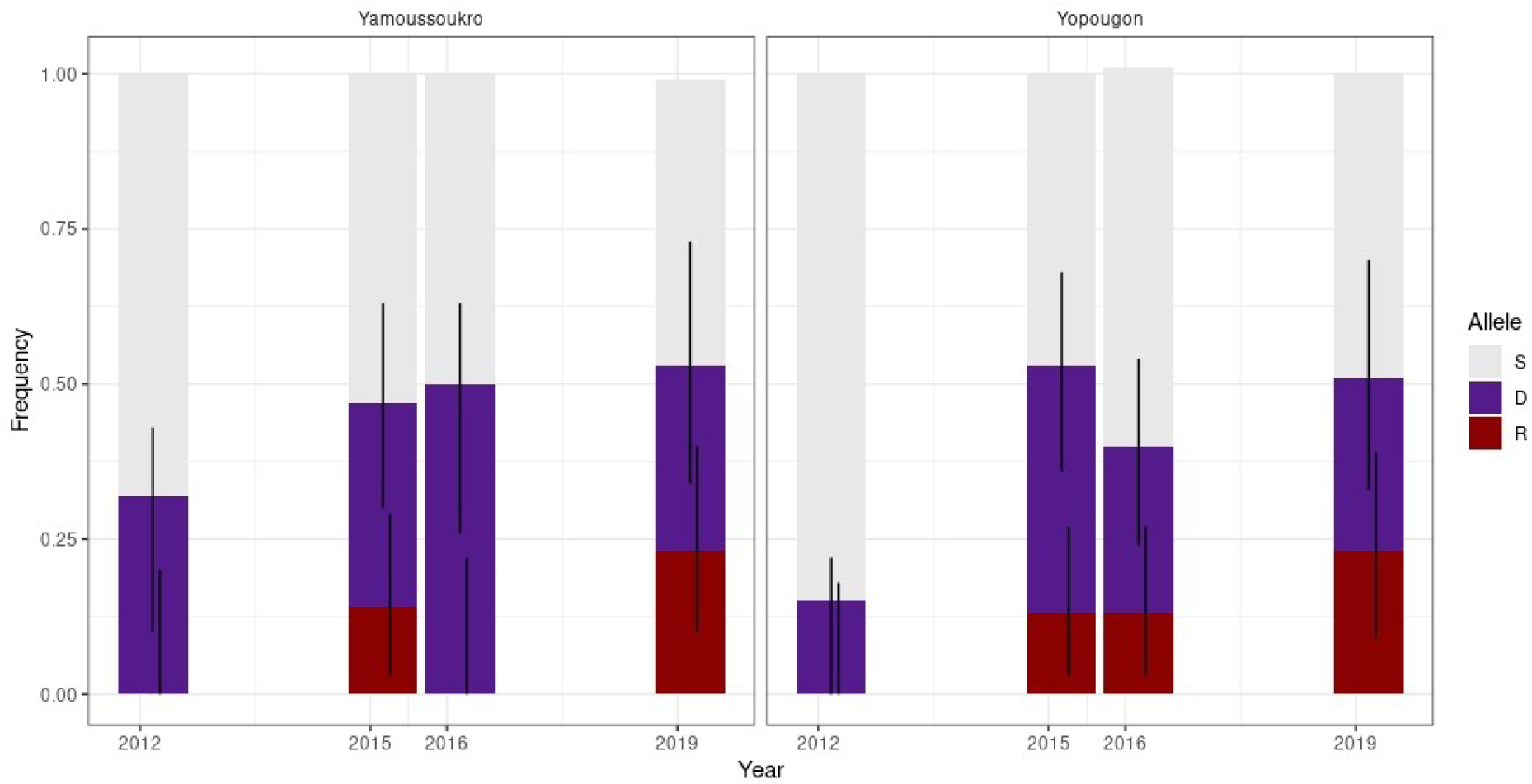
R, S and D allelic frequencies estimated in Yamoussoukro and Yopougon. Data from Assogba *et al*. 2018 (years 2012 to 2016), and from 2019 (this study) were used. We estimated the allelic frequencies considering a 3-allele model (all D alleles are considered together, Tab. 1 Model A) through a maximum likelihood approach (see also *Estimation of duplicated genotype frequencies* in Materials); they are shown in stacked plots with bars corresponding to upper and lower support limits (≈95% confidence intervals).

**Supporting information Table 1.**
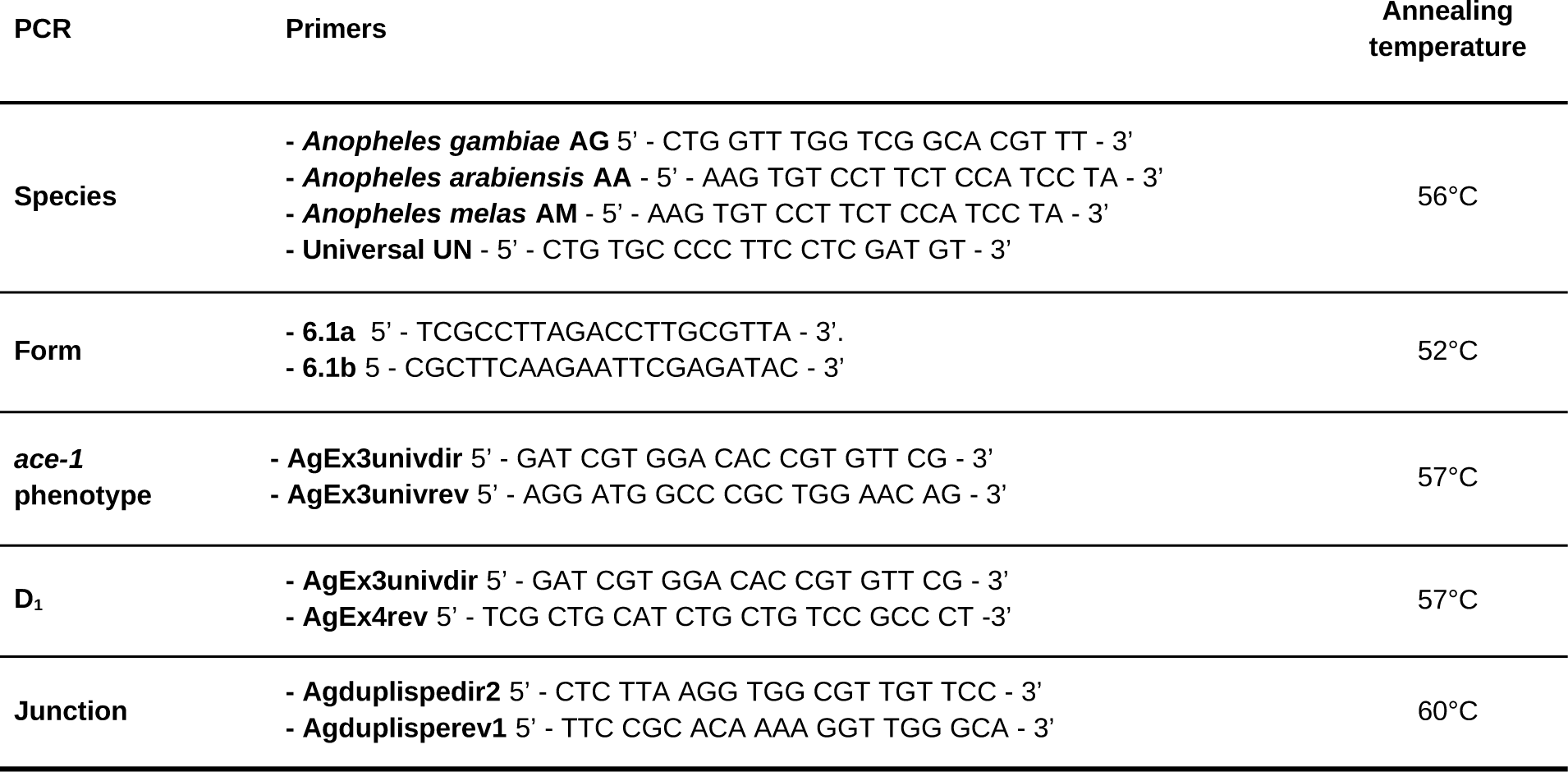
Molecular tests. Various PCR were used for identifying the species (Species) and the molecular form (Form) of the analyzed individuals, as well as their *ace-1* phenotype (*ace-1* phenotype), whether they carry the D_1_ allele (D_1_), and whether they carry a duplicated allele sharing the same junction between amplicons as D_1_ and R^x^ alleles (Junction). The primers used and the reaction characteristics are indicated.

**Supporting Information Table 2.** Sequences dataset from previously published studies. The different sequences used in this study are indicated with their NCBI accession number, the date of collection and locality/country where they were collected, and the corresponding study.

**Supporting information Table 3.**
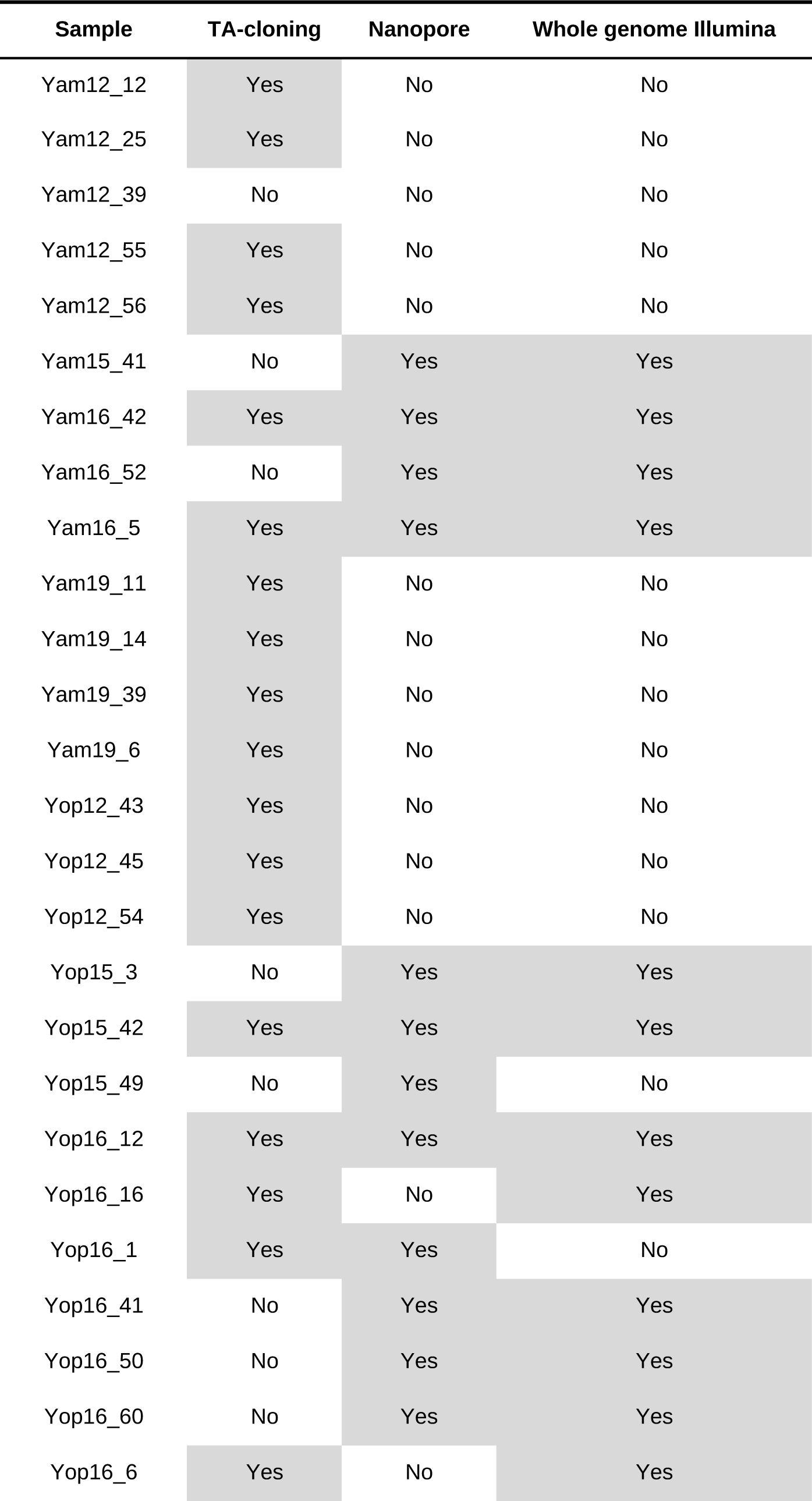
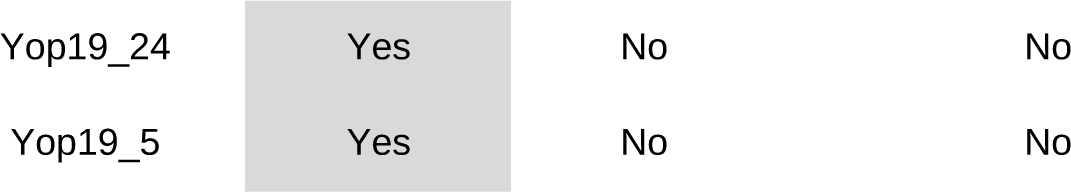
Summary of sequencing and cloning protocol used for each sample of our study. For each individual analyzed in the present study, we summarized here those that were analyzed at the *ace-1* locus, whether cloned and/or sequenced with long-reads (Nanopore), and those whose whole genome was sequenced (Illumina) (see Materials).

| Sample | TA-cloning | Nanopore | Whole genome Illumina |
| --- | --- | --- | --- |
| Yam12_12 | Yes | No | No |
| Yam12_25 | Yes | No | No |
| Yam12_39 | No | No | No |
| Yam12_55 | Yes | No | No |
| Yam12_56 | Yes | No | No |
| Yam15_41 | No | Yes | Yes |
| Yam16_42 | Yes | Yes | Yes |
| Yam16_52 | No | Yes | Yes |
| Yam16_5 | Yes | Yes | Yes |
| Yam19_11 | Yes | No | No |
| Yam19_14 | Yes | No | No |
| Yam19_39 | Yes | No | No |
| Yam19_6 | Yes | No | No |
| Yop12_43 | Yes | No | No |
| Yop12_45 | Yes | No | No |
| Yop12_54 | Yes | No | No |
| Yop15_3 | No | Yes | Yes |
| Yop15_42 | Yes | Yes | Yes |
| Yop15_49 | No | Yes | No |
| Yop16_12 | Yes | Yes | Yes |
| Yop16_16 | Yes | No | Yes |
| Yop16_1 | Yes | Yes | No |
| Yop16_41 | No | Yes | Yes |
| Yop16_50 | No | Yes | Yes |
| Yop16_60 | No | Yes | Yes |
| Yop16_6 | Yes | No | Yes |
| Yop19_24 | Yes | No | No |
| Yop19_5 | Yes | No | No |

**Supporting information Table 4.** Probable genotypes of D-carrying individuals. The probable genotype of the 28 individuals identified as carrying at least one D allele, *i.e.* the triple-peak individuals, has been inferred from the phylogram (Fig. 2) and the genomic analyses, as described in Results. Each individual carried two S copies (S1 and S2, randomly assigned), one being D_i_(S) and the other one being either a single-copy S allele, or another D_j_(S) copy. Each D allele is colored as in Fig. 2.

| Individual | Copy S1 | Copy S2 |
| --- | --- | --- |
| Yam12-12 | D3(S) | S |
| Yam12-25 | D2(S) | S |
| Yam12-39 | S | D2(S) |
| Yam12-55 | D3(S) | S |
| Yam12-56 | S | D2(S) |
| Yam15-41 | D7(S)? | D7(S)? |
| Yam16-42 | D8(S)? | D8(S)? |
| Yam16-5 | D4(S) | S |
| Yam16-52 | S | D4(S) |
| Yam19-11 | D1(S) | S |
| Yam19-14 | D1(S) | S |
| Yam19-39 | D1(S) | S |
| Yam19-6 | D1(S) | S |
| Yop12-43 | S | D1(S) |
| Yop12-45 | S | D5(S) |
| Yop12-54 | S | D1(S) |
| Yop15-3 | D9(S)? | D9(S)? |
| Yop15-42 | D2(S) | S |
| Yop15-49 | S | D6(S) |
| Yop16-1 | D1 | S |
| Yop16-12 | S | D2(S) |
| Yop16-16 | D2(S) | S |
| Yop16-41 | D2(S) | S |
| Yop16-50 | D2(S) | S |
| Yop16-6 | S | D2(S) |
| Yop16-60 | D2(S) | D3(S) |
| Yop19-5 | S | D1(S) |
| Yop19-24 | D2(S) | S |

